# Brain-Cognitive Gaps in relation to Dopamine and Health-related Factors: Insights from AI-Driven Functional Connectome Predictions

**DOI:** 10.1101/2024.11.19.624334

**Authors:** Morteza Esmaeili, Erin Bjørkeli, Robin Pedersen, Farshad Falahati, Jarkko Johansson, Kristin Nordin, Nina Karalija, Lars Bäckman, Lars Nyberg, Alireza Salami

## Abstract

A key question in human neuroscience is to understand how individual differences in brain function relate to cognitive differences. However, the optimal condition of brain function to study between-person differences in cognition remains unclear. While many studies have developed objective biomarkers to accurately predict intelligence and general cognition, consensus on domain-specific markers has not yet emerged. Brain age has been proposed as a potential candidate, but recent research suggests that brain age offers minimal additional information on cognitive decline beyond what chronological age provides, prompting a shift toward approaches focused directly on cognitive prediction. Using a deep learning approach, we evaluated the predictive power of the functional connectome during various states (resting state, movie-watching, and n-back) on episodic memory and working memory performance. Our findings show that while connectomes during task, especially during movie watching, better predict both episodic and working memory, resting state connectomes are equally effective in predicting episodic memory. Furthermore, individuals with a negative brain-cognition gap (where brain predictions underestimate actual performance) exhibited lower physical activity and higher cardiovascular risk compared to those with a positive gap. This shows that knowledge of the brain-cognition gap provides insights into factors contributing to cognitive resilience. Further lower PET-derived measures of dopamine binding were linked to a greater brain-cognition gap, mediated by regional functional variability. Together, our findings highlight the importance of brain state in connectome-based cognitive prediction and introduce the brain cognitive gap as a potentially informative, dopamine-modulated marker of vulnerability to compromise brain function.

## 1. INTRODUCTION

Resting state functional magnetic resonance imaging (fMRI) serves as a tool to map brain function (1, 2). Functional connectivity (FC) at rest, estimated with methods to gauge correlated spontaneous activity between two or more regions, serves as a measure of functional brain integrity (1, 3). Individual differences in FC at rest are associated with differences in various cognitive domains (4–6). Associations of this nature have been reported for several large-scale networks, including the default mode network (DMN) (7–9), the frontoparietal network (10–12), the dorsal attention network (13), and the salience network (14). Moreover, such associations have also been identified in more comprehensive investigations spanning the entire functional brain repertoire of the brain (15, 16). Most previous reports focused on associations, not predictions, utilizing the correlation between FC and behavioral phenotypes, which tend to overfit the data and therefore fail to generalize (17). Proper cross-validation, preferably in an independent sample, is therefore important to assert reliable population-level inferences (18). Recent machine learning-based predictive frameworks offer powerful tools for assessing the predictability of individual behavioral phenotypes based on brain connectivity (19–23). In particular, deep neural networks (DNN) methods have been successfully applied to behavioral and disease prediction (24–26), and were initially expected to outperform other machine learning approaches (27–29). However, this superiority remains debatable, as recent studies have reported comparable performance between DNNs and traditional methods (29, 30). Accordingly, the present study does not aim to benchmark deep learning against traditional machine learning approaches but instead uses a consistent predictive framework to examine how brain state influences the utility of FC for cognitive prediction.

The functional connectome has demonstrated predictive utility regarding trait-like cognitive phenotypes (31–34). The predictive-modeling framework of the functional connectome has been applied to various cognitive domains, including intelligence (35, 36), working memory (WM) (37), visuospatial ability (38), attention (39), creativity (40), as well as personality traits (41). Understanding the patterns contributing to predictions could offer insights into the functional organization underlying cognitive phenotypes, serving as biomarkers indicating current or prospective health conditions (42–44). Moreover, the whole-brain functional connectome acts as a fingerprint with accurate identification of subjects from a large population (45) within the same cognitive state (e.g., rest-to-rest) but also across different states (e.g., rest-to-task). Overall, past research suggests that the functional connectome is relatively robust within individuals, is unique across individuals, and can predict cognitive and personality phenotypes. However, less is known about how the predictive utility of the functional connectome depends on the brain state during which FC is measured.

Despite consensus on the value of resting state functional connectivity for mapping brain function, there is an ongoing debate about whether rest is the optimal brain state for investigating individual differences in neurocognitive function (46–49). A study using data from the Human Connectome Project has shown that resting state fMRI predicts differences in brain activity during various tasks, including social, language, relational, and motor tasks (50). This finding supports the notion that individual differences in neural activation can be predicted from resting state (48). However, results from the same dataset revealed that FC during task outperforms resting state FC in predicting individual differences in fluid intelligence, with FC during task explaining 20 % of the variance compared to just 6 % explained by rest (51). Consistent with this result, previous studies have investigated trait-as well as state-dependent FC, supporting the utility of an integrative approach (11, 49). More recent studies suggested that naturalistic viewing, such as movie-watching, may serve as a happy medium between unconstrained rest and overly-constrained tasks in predicting behavior differences (52, 53). Despite the presence of similar spatiotemporal activity patterns across individuals during movie-watching (54), notable individual differences in activity and functional connectivity (55) persist alongside these idiosyncratic features. This suggests that tasks which align individuals’ functional connectome more closely to an optimal level, neither completely unconstrained like rest nor overly synchronized like a task, also render them easiest to identify (46). Hence, it is plausible that FC during naturalistic paradigms improves sensitivity to predict behavioral differences (52, 56).

The primary objective of this study is to determine how brain state influences the predictive utility of the functional connectome for cognitive performance, using a deep learning framework. Specifically, we test whether functional connectivity derived from different brain states differentially predicts WM and episodic memory (EM), two cores but functionally distinct cognitive domains. WM reflects the capacity to temporarily store and manipulate information and supports higher-order problem-solving, reasoning, and other key components of fluid intelligence (e.g., (57–59)), whereas, EM entails the recollection of specific experiences and events (60), which is regarded as an important element of mind-wandering (61) during resting state. We examine whether these domains are differentially predicted by connectomes derived from resting state, movie watching, and n-back task fMRI.

Past studies have used neuroimaging data, including resting state, to predict brain age (62–64). These studies show that brain age, based on biological phenotypes, and their deviation from the chronological age (known as brain age gap prediction), could serve as a biomarker in characterizing disease risk (64–66). Importantly, an older-appearing brain has been shown to exacerbate physiological and cognitive aging and risk of mortality (63). A recent study demonstrated that while brain age can predict chronological age with high accuracy from MRI, its utility for predicting cognition is limited (67). Specifically, Tetereva and colleagues (2024) (67) showed that brain age strongly tracks chronological age and that, to predict cognition, brain age largely overlapped with chronological age, such that controlling for chronological age eliminated the predictive contribution of brain age. This finding suggests that brain-age models may provide little unique explanatory power for cognitive decline beyond what is already captured by chronological age. Building on this observation and extending the concept of a brain age gap to a brain-cognition gap (BCG, defined as the discrepancy between predicted and observed cognitive performance), we propose that BCG may serve as an informative marker of individual differences. If the brain predicts lower performance than is observed (i.e., a negative BCG), it may be compensating for underlying issues not yet apparent through cognitive assessments. By this view, individuals with negative BCG should be less healthy than those whose brains predict higher cognitive function than their actual performance (i.e., a positive BCG). Our second aim was to extend the concept of brain-age prediction to cognition by introducing BCG. Considering the significance of lifestyle and cardiovascular risk for maintaining healthy brain function (68, 69), we assess whether BCG captures individual differences beyond chronological age and examine whether individuals with positive versus negative BCG differ in lifestyle factors and cardiovascular risk, which are known contributors to brain health.

The third aim of the current study is to investigate the neurobiological underpinnings of BCG by examining the role of dopaminergic (DA) integrity. We test the hypothesis that lower DA receptor availability is associated with increased blood-oxygen-level-dependent (BOLD) signal variability, reduced functional connectome uniqueness, and larger BCG, consistent with DA’s role in modulating neural signal-to-noise ratio (SNR) and network coherence. DA is a vital neuromodulator with critical implications for motor function, reward-seeking behavior, and various higher-order cognitive functions (70–77). Insufficient DA modulation can affect neurocognitive functions detrimentally (71, 76, 78–80). (81, 82) (83, 84) (85). Pharmacological studies have shown that DA depletion increases the variability of the BOLD signal, subsequently leading to less synchronized connectivity within resting state networks (86). Consequently, we expect individuals with inadequate DA levels to exhibit increased regional signal variability, a less unique functional connectome, and greater BCG.

Using data from the DopamiNe, Age, connectoMe, and Cognition (DyNAMiC) study (n =180, 20-79 years, 50% female) (56), we evaluated the predictive power of the functional connectome during resting state, movie-watching, and n-back tasks for two different cognitive domains: episodic memory and working memory. Based on recent research indicating that movie-watching enhances predictability by highlighting key features of FC (52), we hypothesized that FC during movie-watching would outperform FC during rest, and possibly during task, in predicting both cognitive measures. To achieve this objective, we employed a deep neural network approach, a specific subtype of artificial intelligence (AI), to predict cognitive scores from the functional connectome. Deep learning approaches offer a flexible modeling framework capable of capturing complex linear and non-linear associations in high-dimensional data (30) and have been shown to reliably predict intelligence (23, 87). Considering the importance of individual characteristics, such as age, in predicting behavior from FC (34), we conducted external validation of our model, initially derived from an age-heterogeneous sample, in an age-homogeneous sample (from the Cognition, Brain, and Aging (COBRA) study (88)). We subsequently investigated whether individuals with positive brain-cognition prediction gaps differ from those with negative gaps in terms of lifestyle and cardiovascular disease risk factors. Moreover, we tested the hypothesis of whether individuals with lower striatal dopamine D1-like receptor availability (D1DR), the brain’s most abundant DA receptor subtype, have a less distinctive FC pattern (i.e., more regional variability) and, in turn, a larger BCG. Finally, we conducted an external validation of the link between DA and the prediction gap in an independent cohort with estimates of DA D2-like receptor (D2DR) availability (88).

## 2. Results

### 2.1. AI-Driven Predictive Modeling of Cognition Scores from the Functional Connectome

We used fMRI data from the DyNAMiC project, in which each subject underwent scanning during rest, movie-watching, and working memory n-back tasks. These data were parcellated into 273 nodes (264 with 9 additional subcortical nodes) using a previously published whole-brain functional atlas (89). The averaged time series of 273 regions were subsequently correlated to create the FC matrix for each participant and cognitive state (rest, movie-watching, and n-back).

We trained a convolutional neural network, DenselyAttention, derived from DenseNet (90) on FC matrices from each condition (resting state, movie-watching, and n-back) to generate cognition-specific prediction models of two memory domains (EM and WM). Model performance was quantified as the correlation between observed and predicted cognitive performance. Each model was then tested on all three conditions to examine the generalizability of each model across cognitive states. For example, the model trained on the resting state data (orange circles in **Fig. 1**) was used to predict EM scores using the test dataset derived from resting state (**Fig. 1a)**, movie-watching, and n-back. Significance of our main predictions was assessed via linear correlations, and uncorrected *p*-values are presented in **Tables 1–2**.

**Table 1.**
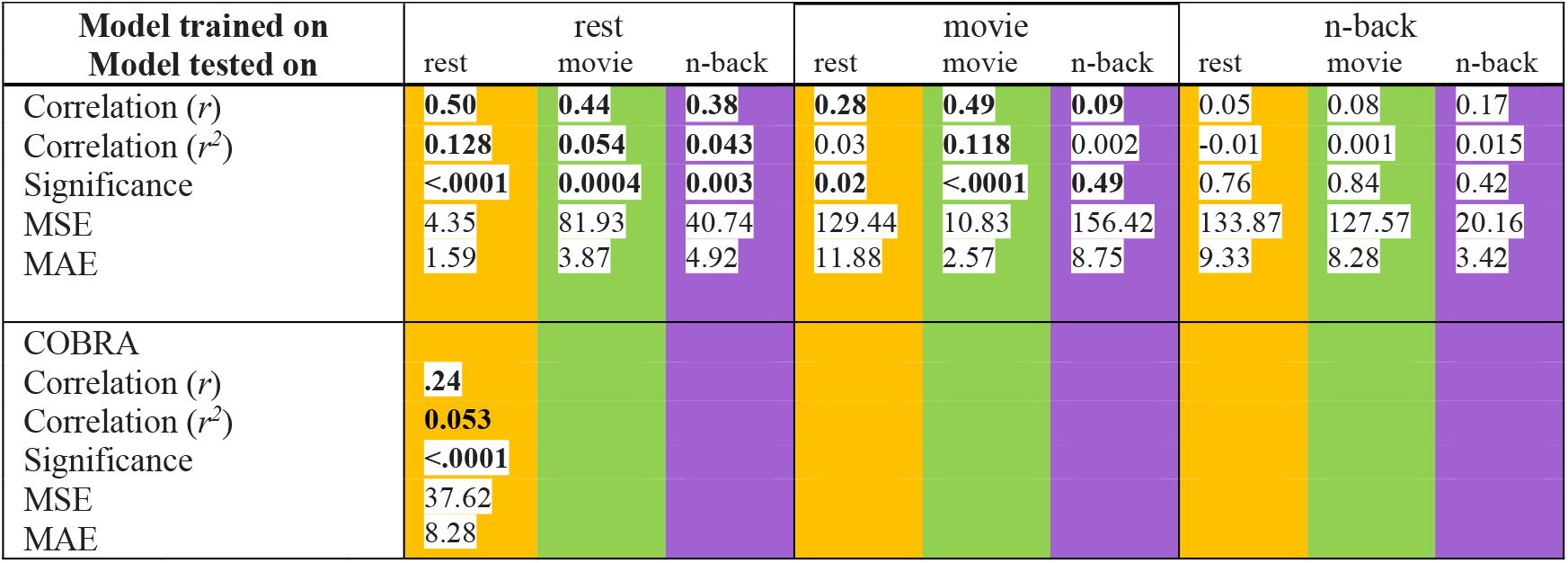
Correlation results for episodic memory (EM) score predictions.

**Figure 1.**
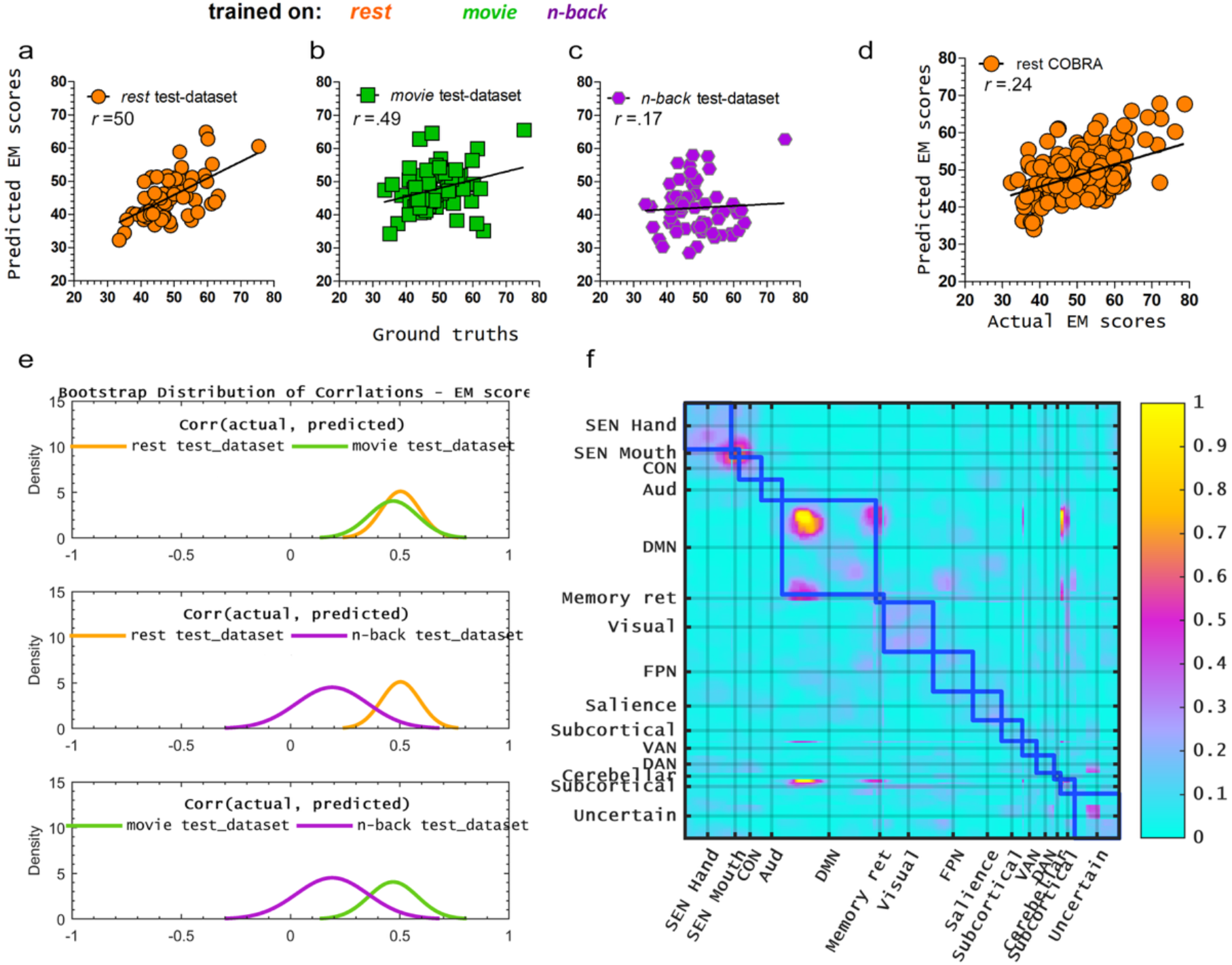
Model trained on functional connectivity maps acquired at rest predicts (**a**) episodic memory (EM) of the test dataset. The models trained on movie-watching (**b**) but not n-back (**c**) datasets predicted EM scores of the test dataset. **Table 1** summarizes the p values, correlations, mean square error (MSE), and mean absolute error (MAE) for each model. Test datasets were obtained from the same cohort for rest, movie-watching, and n-back. The winning model trained on EM at rest was evaluated on the external COBRA cohort and yielded a significant prediction of EM scores (**d**). Bootstrap distributions of correlations between predicted and actual EM scores indicated no significant difference in the predictive power of EM between models trained on resting state and movie-watching data (**e**). Additionally, the bootstrap distribution revealed that models trained on resting state and movie-watching data yielded higher correlations than those trained on n-back data (**e**). Visualization of features contributing to the successful prediction of EM at rest (**f**). A grad-CAM-derived saliency map displays the features that contributed to the model’s predictions. The hot spots overlaid on the FC map demonstrate noticeable cross-correlation contributions in “default mode” (DMN) regions. Another important feature visualized by Grad-CAM includes off-diagonal hot spots reflecting inter-connections of the DMN – “subcortical” node.

### 2.2. Resting state and movie-watching models outperform n-back in episodic-memory prediction, with resting state offering the best generalizability

We first started with all cases in which congruent conditions were used for model building and prediction. Only models derived from the rest and movie-watching datasets yielded significant predictions of EM (**Figs. 1a-b** and **Table 1**), with resting state yielding the best-performing model (*r* =0.50, *p* <0.0001), followed by movie-watching (*r* =0.49, *p* <0.0001) (**Table 1**). While there was no significant difference between resting state and movie watching in predicting episodic memory (Δr =0.071, with a 95% confidence interval of [-0.097, 0.261], **Fig. 1e**), both models yielded a markedly better EM prediction than n-back (rest vs. n-back: Δr =0.333, with a 95% confidence interval of [0.054, 0.572]; movie vs. n-back: Δr=0.316, with a 95% confidence interval of [0.015, 0.619], **Fig. 1e**). Thus, the two models outperformed the n-back model (*p* <0.05, bootstrap test), indicating a significant improvement. To test the generalizability of these models, two types of validation analyses were performed: cross-condition and cross-data set. In the cross-condition analysis, models trained on one condition (e.g., rest) were tested on an incongruent condition (e.g., movie-watching, n-back; **Table 1**). Interestingly, the model trained on resting state significantly predicted EM when tested on movie-watching (*r* =0.44, *p* <0.001) and n-back (*r* =0.38, *p* =0.003) conditions. In contrast, models trained on movie-watching or n-back could not be generalized to other conditions, unable to significantly predict EM (*p*’s >.1), except for significant generalizability from the movie to the rest condition (**Table 1**).

In a cross-dataset validation analysis, the best-performing model from the age-heterogeneous DyNAMiC dataset was tested on the corresponding condition in an age-homogeneous cohort from the COBRA dataset. By doing this, we found that the resting state model derived from DyNAMiC significantly predicted EM performance in the COBRA dataset (*r* =0.24, *p* <0.0001). Next, we aimed to delineate the relative contributions of different brain regions for the best-performing model, the model trained on the “resting state data” in predicting episodic memory. Utilizing the Grad-CAM algorithm, saliency maps were generated for the 120 FC matrices used during training of the winning model. An averaged and interpolated description of all saliency maps is depicted in **Figure 1f**. The saliency map highlights specific edges, especially within the default-mode network, edges between the default-mode network and subcortical areas, and edges between the default-mode and the cerebellar network. These edges, indicated by a salience intensity of ≥0.5, exert a significant influence on the model (**Fig. 1f**).

### 2.3. Movie-watching and n-back models outperform resting state in working-memory predictions, with movie-watching offering the best generalizability

We next investigated whether the superiority of resting state in predicting EM was unique to this domain, considering that previous research demonstrated the advantages of task-based fMRI and naturalistic viewing in predicting fluid intelligence (91). To do so, we compared the predictive power of different states for WM, which is shown to be more directly associated with fluid intelligence compared to EM.

The model derived from the resting state failed to predict and generalize regarding WM (*p*’s >0.10; **Fig. 2a** and **Table 2**). By contrast, models trained on movie-watching and n-back yielded significant predictions of WM (**Figs. 2b-c** and **Table 2**), with the movie-watching model emerging as the best-performing model (*r* =0.57, *p* <0.0001) followed by n-back (*r*^2^ =0.47, *p <*0.0001). While there was no significant difference between movie-watching and n-back in predicting WM (Δr =0.026, with a 95% confidence interval of [0.001, 0.052], **Fig. 2e)**, these models yielded better WM prediction than resting state (Δr =0.517, with a 95% confidence interval [0.373, 0.662], **Fig. 2e**).

**Table 2.**
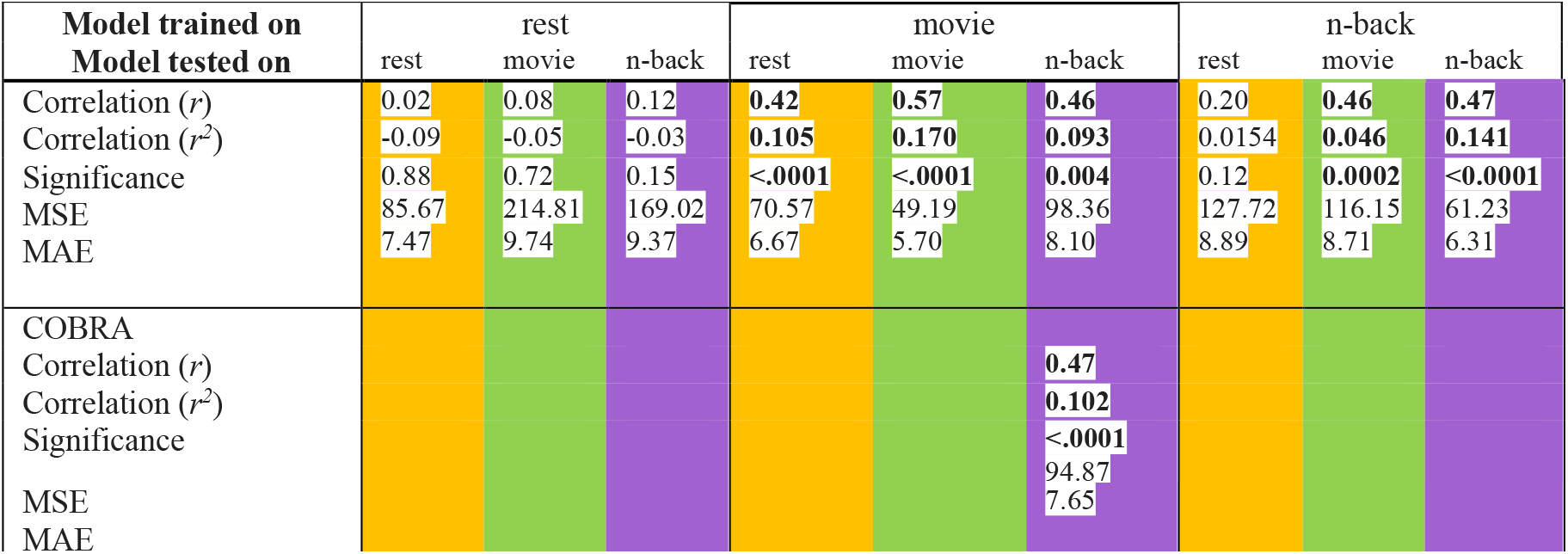
Correlation results for working memory (WM) score predictions.

**Figure 2.**
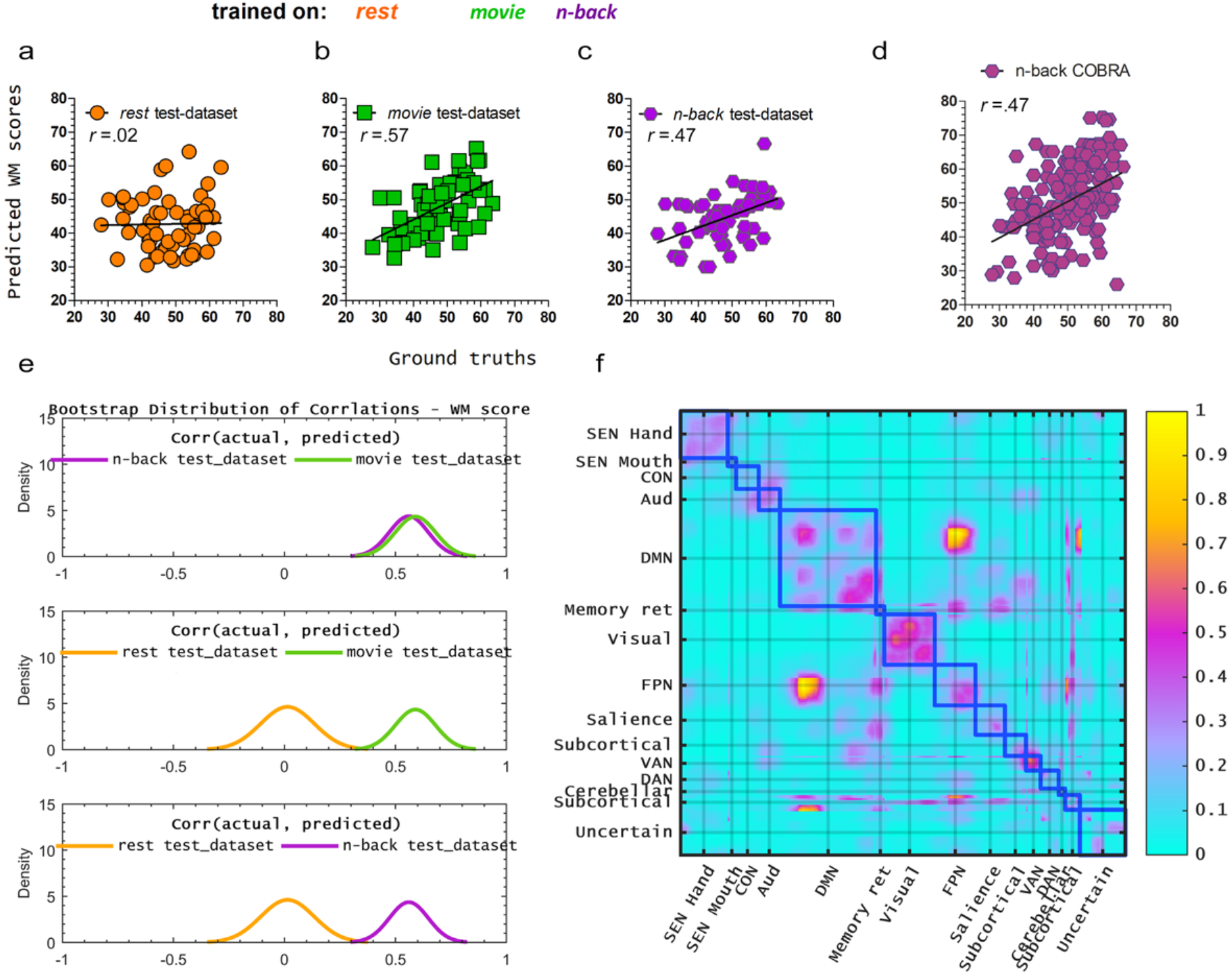
Model trained on FC maps acquired at rest did not significantly predict (**a**) the working memory (WM) in the test dataset. The model trained on the movie-watching dataset yielded the best-performing model in predicting WM (**b**) while the model trained on the n-back dataset (**c**) was the second-best model. **Table 2** summarizes the p-values, correlation power, MSE, and MAE for each model. (**d**) Results of cross-dataset validation, where the best-performing model in the DyNAMiC dataset (i.e., movie-watching) was applied to predict WM to the COBRA dataset. However, since COBRA does not include a movie-watching paradigm, we applied the model to the n-back task in COBRA (**Table 2**). Bootstrap distributions of correlations between predicted and actual WM scores showed no significant difference in predictive power between models trained on movie-watching and n-back data (**e**). The bootstrap distribution revealed that models trained on movie-watching and n-back data exhibited higher correlations than those trained on resting state data (**e**). The Grad-CAM-derived saliency map highlights dominant features in the FC maps that contributed to the model’s predictions (**f**). The hot spots overlaid on an FC map demonstrate noticeable cross-correlation contributions in the “VAN”, “visual”, and to a lesser degree (<0.5) “DMN”. Other important features visualized by Grad-CAM include off-diagonal hot spots reflecting inter-connections of the “DMN” – with “FPN, Fronto-parietal Task Control”, “Subcortical”, and “Cerebellar”; “Cerebellar” – “FPN” node.

In cross-condition validation (**Table 2**), the movie-watching model yielded a significant WM prediction during both resting state (*r* =0.42, *p* <0.0001) and n-back (*r* =0.46, *p* =0.004). The model derived from n-back yielded a significant WM prediction during movie-watching (*r* =0.46, *p* =0.0002), but not during resting state (*r* =0.20, *p* =0.12).

For cross-dataset validation, the best-performing model in the DyNAMiC dataset (i.e., movie-watching) was applied to predict WM in the COBRA dataset. However, since COBRA does not include a movie-watching paradigm, we applied the model to the n-back task in COBRA. This approach revealed that the model from the DyNAMiC movie-watching condition yielded a significant WM prediction in the COBRA n-back task (*r* =0.47, *p* <0.0001).

Overall, our results suggest that movie-watching-based WM predictions generalize across different cognitive states and datasets. This finding could be further replicated using a different functional parcellation (**Figs. S1-S2 and Tables S1-S2**).

The Grad-CAM algorithm generated saliency maps of 120 FC maps from the DyNAMiC dataset employed for training. **Figure 2f** depicts an average of all Grad-CAM generated maps. The saliency map unveils that certain edges, specifically within network connectivity of task-positive regions such as the frontoparietal task control network, dorsal/ventral attention, visual, and subcortical networks, as well as between-network connectivity. FC between task-positive dorsal and ventral attention networks, and between the DMN and the fronto-parietal network, contributed to the best-performing model derived from the movie-watching dataset. Applying two different parcellation methods (89, 92) to the DyNAMiC data indicated that parcellation resolution does not significantly impact model performance (see **Figs. S1-S2 and Tables S1-S2**).

### 2.4. The brain-cognition gap is related to physical activity levels and cardiovascular risk factors

Given the importance of lifestyle and cardiovascular health for maintaining healthy brain function (68, 69, 93), we examined whether individuals with positive versus negative prediction gaps differed in physical activity habits, education, and Framingham cardiovascular disease (CVD) risk score. Our primary focus was on EM predictions derived from resting state data, as this was the common condition across the DyNAMiC and COBRA datasets. We computed the difference between predicted and observed EM scores to generate BCG. A positive BCG indicates that an individual’s brain predicted better-than-observed EM performance, whereas a negative BCG indicates more compromised brains relative to actual performance.

In the test sample of the DyNAMiC data (n =60) and the entire COBRA sample (n =177), we found that individuals with a negative BCG exhibited lower levels of physical activity and higher CVD risk scores compared to those with positive gaps (**Fig. 3**). Confirmatory analysis with continuous variables revealed positive relationships between GAP and physical activity (DyNAMiC: *r* (57) =0.40, *p* =0.001; COBRA: *r*(166) =0.17, *p*=0.03) and negative relationships between GAP and CVD risk score (DyNAMiC: *r*(57) =–0.27, *p* =0.03; COBRA: *r*(172)= –0.10, *p* =0.40). Moreover, individuals with negative BCG were less educated compared to those with positive BCG in the DyNAMiC dataset.

**Figure 3.**
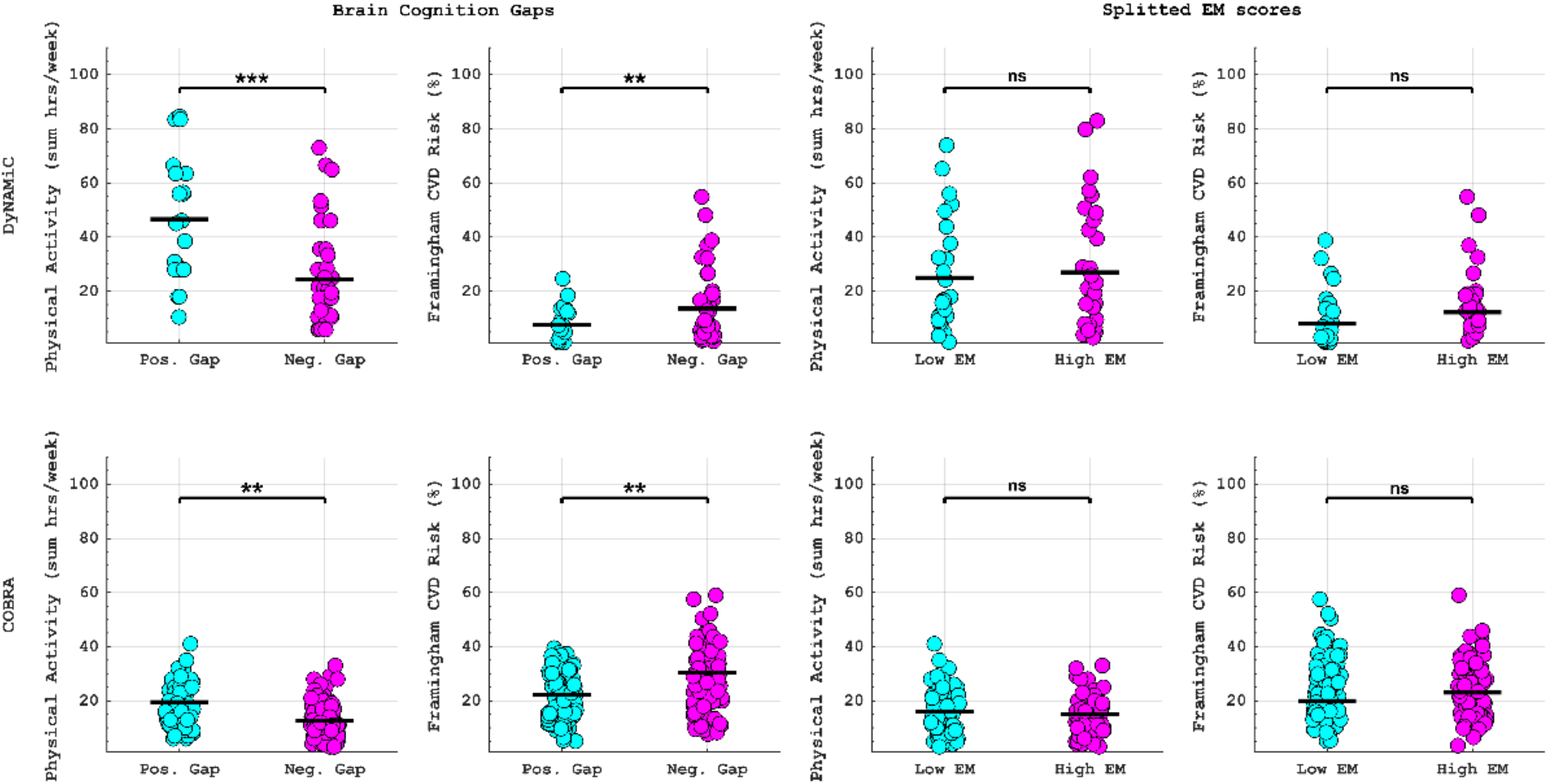
The plots compare physical activity scores (total hours per week), CVD risk, between two groups with positive and negative BAG as well as high and low memory performance in the DyNAMiC (**top row**) and COBRA (**bottom row**) datasets. *, **, and *** denote p <0.05, p <0.01, and p <0.001 respectively.

To test whether cognition on its own is related to physical activity and CVD score, we conducted a median split on EM and compared physical activity and cardiovascular risk score across the two groups. In contrast to the findings related to BCG, we found no significant difference in the level of physical activity (t(58) =-0.59, *p* =0.56) or cardiovascular risk score (t(58) =1.64, *p* =0.11) between high and low EM performers (**Fig. 3**). These results suggest that BCG may provide additional information, beyond cognitive measures, regarding factors that contribute to cognitive resilience.

### 2.5. Dopamine D1 and D2 receptor availability are associated with brain-cognition gaps

Given that BCG may partly reflect variability in neural signal, one plausible neurobiological factor contributing to BCG is dopaminergic integrity. We hypothesized that inadequate DA levels might be related to increased neural signal-to-noise ratio, thereby resulting in a less unique functional connectome, consequently leading to a greater prediction gap. We therefore initially investigated the relationship between DA receptor levels and predictive gaps across different types of DA receptors in the DyNAMiC and COBRA samples.

In the DyNAMiC sample, we found a significant correlation between striatal D1DR and prediction BCG (in those with a positive gap: *r* = –0.49, *p* =0.03; negative gap: *r* =0.40, *p* =0.01) (**Fig. 4a**), suggesting that lower D1DR is associated with greater BCG.

**Figure 4.**
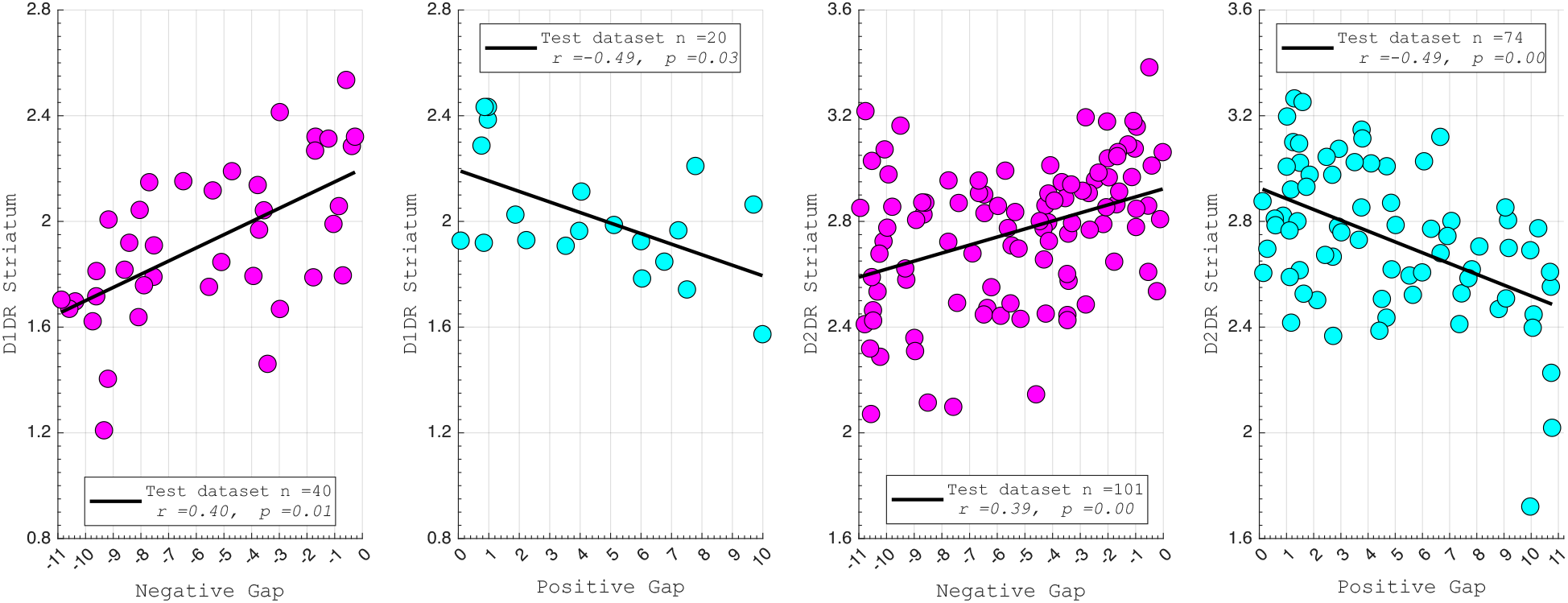
Relationship between the gap measured from predicted and actual EM scores and dopamine D1 receptor (**a**). Negative gaps indicate that the predicted EM score was lower than actual EM scores, while the positive gaps indicate more higher predicted scores than actual EM scores. Partial correlation analysis showed a significant correlation between D1 receptor values and the measured negative and positive gaps. Relationship between the gap measured from predicted and actual EM scores and dopamine D2 receptor (**b**). Negative gaps indicate that the predicted EM score was lower than actual EM scores, while positive gaps indicate more higher predicted scores than actual EM scores. Partial correlation analysis showed a significant link of D2 receptor values to negative gaps and positive gaps.

We replicated our finding in the COBRA sample using D2-like receptor availability (D2DR), revealing a significant relationship between striatal D2DR and prediction gap (in those with a positive gap: *r* = –0.49, *p* =0.001; negative gap: *r* =0.39, *p* =0.004) (**Fig. 4b**). Our findings provide support the view that lower D1DR/D2DR is associated with larger brain-cognition prediction gaps.

Both D1DR and D2DR availability in the striatum were associated with BCG, such that lower dopamine receptor availability was linked to a greater BCG. However, these associations varied by region. For D1DR, significant correlations with BCG were observed in the caudate (positive gap: *r* = –0.37, *p* =0.02; negative gap: *r* = 0.37, *p* =0.02) and putamen (positive gap: *r* = – 0.53, *p* =0.02; negative gap: *r* =0.34, *p* =0.03), but not in the nucleus accumbens (positive gap: *r* = –0.25, *p* = 0.31; negative gap: *r* =0.07, *p* =0.69) or the DLPFC (positive gap: *r* = – 0.30, *p* =0.21; negative gap: *r* =0.21, *p* =0.21). For D2DR, both caudate (positive gap: *r* = – 0.34, *p* =0.004; negative gap: *r* =0.36, *p* =0.0003) and putamen (positive gap: *r* = – 0.37, *p* =0.002; negative gap: *r* =0.22, *p* =0.03) showed significant associations with BCG.

### 2.6. Regional variability mediates the direct impact of dopamine on brain-cognition gaps

We showed that dopamine is associated with BCG. To evaluate whether functional variability mediated this relationship, we conducted additional mediation analyses. We computed BOLD signal entropy, which estimates within-region signal variability during resting state (86), that was averaged across striatal regions (left caudate: MNI coordinate ≈ [–12, 12, 6], right caudate: MNI coordinate ≈ [10, 14, 2], left putamen: MNI coordinates ≈ [–24, 6, 4], right putamen: MNI coordinates ≈ [29, 1, 4]), as we expected that reduced DA may primarily impact the local functional dynamics.

In the DyNAMiC sample, within the group exhibiting a negative BCG, we observed a negative association between striatal D1DR and entropy (*r* =–0.33, *p* =0.04) as well as a negative association between entropy and BCG (*r* =–0.36, *p* =0.03). Importantly, we observed a significant indirect effect of D1DR on the gap mediated by entropy, *β* =2.41, 95% CI [0.89, 4.51], *p* <0.0001, explaining 56.81% of the total effect of D1DR on the gap. The direct effect of D1DR, however, was not significant, *β* =1.83, 95% CI [–0.86, 3.79], *p* =0.19.

Similarly, in the group with a positive BCG, entropy was negatively associated with D1DR (*r* =–0.56, *p* =0.01) and positively associated with BCG (*r* =0.47, *p* =0.04). Importantly, an indirect effect of D1DR on BCG through entropy was observed, *β* =–6.78, 95% CI [–11.26, –3.05], *p* <.0001, accounting for 89.41% of the D1DR effect on the gap. The direct effect of D1DR was again non-significant, *β* =–0.803, 95% CI [–0.48, 2.27], *p* =0.26. These findings suggest that lower D1DR levels contribute to increased signal variability, which in turn leads to reduced specificity of FC and, consequently, a larger BCG.

In the COBRA sample, within the group with a negative BCG, we observed a negative association between striatal D2DR and entropy (*r* = –0.22, *p* =0.03) as well as a negative association between entropy and BCG (*r* = –0.27, *p* =0.007). In the group with the positive BCG, entropy was negatively associated with D2DR (*r* = –0.26, *p* =0.03) and positively associated with BCG (*r* =0.25, *p* =0.03). Moreover, we detected a significant indirect effect of D2DR on both the negative and positive gap groups through entropy. For the negative gap, *β* =2.18, 95% CI [0.01, 4.25], *p* =0.04, accounting for 63.43% of the D2DR effect on the gap; and for the positive gap, *β* =–2.18, 95% CI [–3.87, –0.04], *p* =0.04, accounting for 61.49% of the D2DR effect on the gap. Similar to the results reported for D1DR, these findings suggest that lower D2DR levels contribute to increased signal variability, which in turn may lead to reduced specificity of FC and, consequently, a larger BCG.

## 3. DISCUSSION

Using deep learning models, we examined the predictive power of the functional connectome during various states (resting state, movie-watching, and n-back) on two different cognitive domains (EM and WM). Both rest and movie-watching states yielded significant predictions of EM, with the model derived from resting state generalized across states and datasets. Differences between the DyNAMiC and COBRA datasets make cross-dataset prediction a harder problem, as the age ranges of samples significantly vary, and prior studies highlight the importance of individual characteristics like age in predicting behavior from FC (34). In line with this, model performance decreased when predicting EM in the COBRA sample, whereas prediction of WM remained largely unchanged. Thus, validation outcomes suggest that the models, particularly those predicting WM, show robustness across datasets, whereas the reduced EM performance highlights potential data-specific influences that limit generalizability. The saliency map generated from the final layer of the deep learning model indicates that certain edges within DMN, and between DMN and the subcortical network contributed significantly to the prediction. Building on a recent finding by Kurkela and Ritchy (94), our finding reveals that a portion of the known-memory subnetwork within the DMN, as well as a whole-brain multivariate pattern which notably encompasses interactions of the DMN with other networks, such as the subcortical network, made a more substantial contribution to prediction. Importantly, this prediction generalizes across conditions and datasets, suggesting that features derived from resting state FC serve as a relatively stable marker of individual differences in EM, though with reduced strength in COBRA. While such generalization is partly facilitated by the similarity of functional connectivity across states, it is not a trivial outcome. For instance, the model trained on movie-watching data generalized to EM prediction during rest but failed to do so for the n-back condition, even though movie-watching and n-back connectivity patterns are themselves highly correlated. This indicates that successful generalization depends not only on shared variance across states but also on the cognitive processes most relevant to the target behavior.

Our findings are in contrast to recent work suggesting that task paradigms, in general, and movie-watching, in particular, outperform resting state data in predicting cognitive performance (51, 52, 95). While previous studies have often demonstrated a superiority of task and naturalistic viewing over resting state in predicting fluid intelligence or WM (51, 52), there are fewer reports of FC predicting EM (e.g., (96, 97)), and, to our knowledge, no study has compared rest and movie-watching. While we acknowledge that the resting state represents a complex amalgamation of cognitive, emotional, and perceptual processes (98), the good prediction power of the resting state may arise from the presence of mind wandering during rest, which is strongly related to EM (99, 100). EM plays a crucial role in generating mental content during mind wandering, especially episodes characterized by distinct times and locations (61, 101).

In contrast to the EM prediction, both n-back and movie-watching connectomes yielded significant predictions of offline WM performance. Importantly, the models derived from movie-watching and n-back outperformed the resting state in WM prediction. These differences in model accuracy when predicting the same target behaviors (i.e., WM) suggest the presence of trait-state interactions. Specifically, movies and n-back enhance individual differences in WM-relevant connections. Indeed, we found that several WM-related networks (102–105), including the fronto-parietal, the salience, and the dorsal/ventral attention networks, contribute to prediction. Additionally, previous research showed that movie-watching alters the propagation of activity across cortical pathways (106), particularly within and between regions involved in audiovisual processing and attention. These alterations lead to a less segregated and more integrated network organization (107). Similarly, the n-back task has been associated with increased integration of task-positive cortico-cortical connectivity (105, 108) and striato-cortical connectivity (103). Our findings also suggest that certain task contexts strike an optimal balance between reducing neural variability and maintaining sufficient richness to capture individual differences. Prior work shows that task states quench neural variability, leading to a more reliable and predictable neural signal (109). In this context, movie watching may represent such a sweet spot constraining neural dynamics through shared audiovisual stimulation, while simultaneously engaging a broad range of cognitive processes that preserve individual differences. Taken together, our results confirm previous findings that movie watching is a suitable condition for studying individual differences across various cognitive domains. Nonetheless, if a movie-watching paradigm is not feasible/ available, resting state still provides a robust means of studying individual differences, particularly in self-referential domains, such as EM.

Our study used a deep neural network architecture that features dense connections and incorporates an attentional mechanism. While our findings demonstrate that a deep learning framework can provide reasonable predictive accuracy, it is important to note that other machine learning approaches (e.g., tree-based models) may offer comparable predictive power, as suggested by prior benchmarking work (29, 30). Our study explicitly compares predictive power across different cognitive states (rest, movie watching, n-back) to identify the states that best capture individual differences across domains. The relative performance of deep learning and other non-linear approaches depends on multiple factors, including sample size, model architecture, feature representation, and domain-specific characteristics of the prediction target. In this context, deep learning was employed as a flexible framework capable of modeling high-dimensional functional connectivity patterns across cognitive states, rather than as a claim of inherent methodological superiority. Thus, our goal was not to propose a universally superior prediction model, but rather to test how brain state influences predictive utility for WM and EM using a deep learning approach.

We found a significant link between BCG, lifestyle, and risk factors for vascular disease, such that individuals with a negative BCG exhibited lower levels of physical activity and higher cardiovascular risk scores compared to those with a positive BCG. This Pattern was observed in both the age-heterogeneous DyNAMiC sample and the age-homogeneous COBRA sample, although the effect sizes were smaller in COBRA This attenuation may reflect differences in sample composition, age range, and assessment of lifestyle-related variables across cohorts. BCG could serve as a potential biomarker for identifying individuals at risk (e.g., individuals with a negative gap). Previous studies suggest that the brain age prediction gap is associated with cognitive aging (64), some aspects of physiological aging (63), as well as aging in other organs (110) and even mortality in older age (63). However, a recent study revealed that brain age accounts for only a small portion of cognitive decline compared to chronological age (67), suggesting that cognitive prediction might be more informative. Our findings build upon this concept by extending the BCG to behavioral variables, demonstrating that the BCG could provide insights regarding physical activity status, education, and cardiovascular risk – key factors contributing to cognitive reserve (111–114). Note that the association with education was significant only in the DyNAMiC sample and did not reach significance in the COBRA dataset. An important caveat is that BCG can also be conceptualized as an error metric, like mean absolute error or mean square error, reflecting the extent to which models trained in one sample generalize to another. From this perspective, a larger gap may not only indicate individual differences related to resilience factors and dopaminergic function but also reduced model fit or generalizability across datasets. Thus, BCG likely reflects a combination of meaningful biological variability and methodological variance.

Critically, we found that D1DRs and D2DRs were strongly associated with the BCG, such that lower dopamine receptor levels were associated with greater gaps. More specifically, in two independent samples, we discovered greater correspondence (i.e., near zero in BCG) between brain function and cognition in individuals with higher D1DR/D2DR, whereas lower correspondence (i.e., significantly different BCG from zero) was found in individuals with lower D1DR/D2DR availability. Previous computational models proposed that DA modulates neuronal gain, which improves SNR in neural processing, contributing to more coherent activity across large-scale networks (e.g., balanced integration and segregation (85)). Past studies also showed that lower D1DR contributed to more BOLD variability in the subcortical area (83) and less functional segregation of the striatum (115) and the large-scale networks in aging (85), possibly due to increased noise (lower SNR). In support of this notion, we found for the first time that regional variability, estimated using entropy, mediated the impact of DA on BCG. Although the cross-sectional nature of our data warrants caution, this novel finding suggests that lower DA integrity relates to BOLD variability, which in turn is associated with a larger BCG.

An important caveat is that D1DR and D2DR availability do not provide a direct measure of dopamine signaling. Instead, it reflects receptor availability, which interacts with endogenous dopamine in a complex manner. PET measures of D1R and D2R availability reflect the density of unoccupied dopamine receptors and the degree to which endogenous dopamine competes with radioligand binding. D2R binding potential is sensitive to competition from synaptic dopamine, such that higher ambient dopamine generally reduces tracer binding; D1R binding, however, is less affected by endogenous dopamine under physiological conditions, reflecting more directly receptor expression levels. Previous studies demonstrated a significant association between D2R availability and dopamine synthesis capacity measured by FMT (116, 117), suggesting that postsynaptic receptor markers may, under certain conditions, serve as a proxy for dopaminergic signaling. Developmental factors, such as the number of dopamine-producing neurons innervating the striatum, may further influence the structural and functional relationship between pre- and post-synaptic markers. By contrast, smaller studies have reported non-significant (118, 119) or negative (120) associations, although these studies relied on [18F]FDOPA, which is considered a less precise index of dopamine synthesis than FMT. Taken together, these reports indicate that the relationship between pre- and post-synaptic markers is complex and not necessarily linear. Accordingly, our observation that lower receptor availability is associated with greater neural variability should not be interpreted as direct evidence of weaker dopaminergic signaling, but rather as reflecting the interplay between receptor density and endogenous dopamine occupancy, particularly in the case of D2DR.

Finally, we did not directly compare BCG and brain-age gap (BAG). While our focus was to investigate whether the BCG provides information about factors contributing to cognitive resilience, we acknowledge that benchmarking BCG against the brain-age gap in predicting lifestyle and vascular risk factors would be valuable. As shown in **Figure S4**, we did not observe significant associations between BAG and these factors, suggesting that BCG may be more sensitive to aspects of cognitive resilience. However, addressing this question lies beyond the scope of the present study, and future work should systematically compare these approaches. Our objective was not to determine whether BCG outperforms BAG, but rather to assess whether BCG provides information beyond cognitive performance itself. We also note that introducing BAG raises additional methodological questions, such as which cognitive state (rest, movie-watching, or n-back) is most appropriate for estimating biological age, warranting a dedicated investigation. Finally, we acknowledge that our main and validation samples are moderate in size for deep learning, which constrains statistical power and generalizability. Although external validation, early stopping, dropout, and regularization help mitigate overfitting, larger samples will be needed in future work to fully establish the robustness of these predictive models.

In summary, our findings reveal that while tasks like movie-watching predict both episodic and working memory, there are features during rest that can effectively predict internally oriented mind-wandering-type tasks, such as EM. Additionally, individuals whose brains predict poorer cognitive performance (i.e., negative gap) exhibit lower physical activity and higher cardiovascular risk compared to those whose brains predict higher cognitive function than their actual performance (i.e., positive gap). This finding suggests that our prediction model offers a potential marker to identify individuals at risk of compromised brain maintenance. Furthermore, individuals with lower DA showed less accurate cognitive prediction (larger BCG) due to increased BOLD variability and less unique and cohesive FC.

## 4. MATERIALS AND METHODS

### 4.1. Participants

All participants provided written informed consent, and studies were conducted in accordance with the Declaration of Helsinki and approved by the Regional Ethical Board and the local Radiation Safety Committee (reference numbers: 2012-57-31M; 2017-404-32M).

This study used data from DyNAMiC (56), which is a longitudinal study with a focus on changes in the brain connectome and the D1DR system. At baseline,180 participants (20-79 years, 50% female) across the adult lifespan underwent all tests between 2017 and 2020 (56) (**Fig. 5**). Rigorous exclusion criteria were used to recruit a sample without neurological conditions and medical treatments affecting brain functioning and cognition. Exclusion criteria included brain injury or neurological disorder, dementia, neurodevelopmental disorder, psychiatric diagnosis, psychopharmacological treatment, history of severe head trauma, substance abuse or dependence, and illicit drug use. Individuals with other chronic or severe medical conditions (e.g., cancer, diabetes, and Parkinson’s disease) were also excluded. Here, we only use data from the baseline measurement.

**Figure 5.**
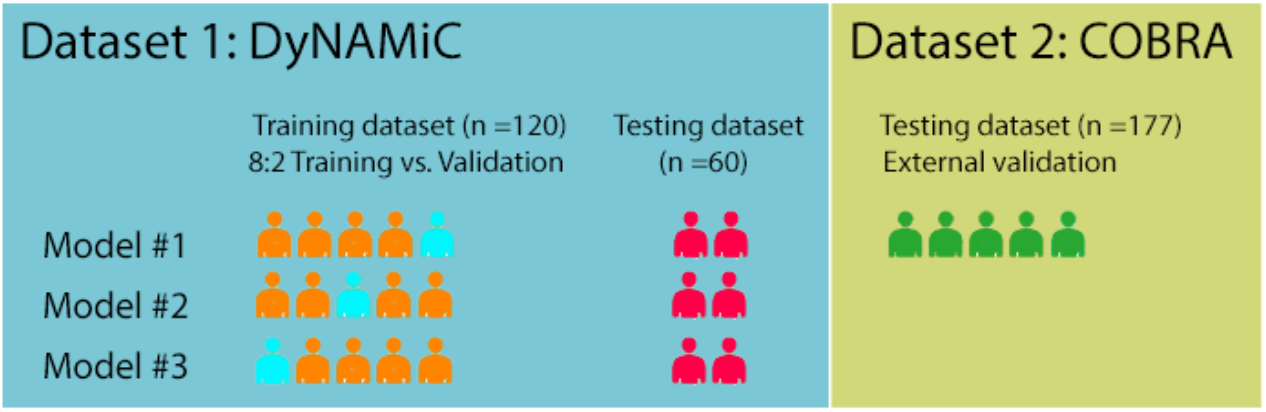
Overview of the experimental procedure and the use of datasets. We used a 3-fold within-sample (DyNAMiC) cross-validation where we trained our model on 120 subjects (8:2; 80% training:20% validation during training) and tested it in a separate sample of 60 subjects. The winning within-sample model was used for between-sample (COBRA) external validation.

We used a separate sample as a testing dataset from the COBRA study (88) (**Fig. 5**). COBRA is a longitudinal aging study in which 181 healthy individuals (64-68 years, 45% female) underwent baseline assessments of the brain, cognition, health, and lifestyle during 2012–2014 (88). Exclusion criteria at baseline included traumatic brain injury, stroke, dementia, intellectual disability, epilepsy, psychiatric and neurological disorders, diabetes, and cancer medications, severe visual or auditory impairment, claustrophobia, and poor Swedish language skills. In the current study, we used data from 177 subjects from COBRA who underwent both MRI and PET examinations at baseline (80, 88, 102, 103, 121).

### 4.2. Cognitive Measures

The same cognitive test battery was used in DyNAMiC and COBRA (56, 88) (**Fig. S1**) and assessed two cognitive domains: episodic memory (EM) and working memory (WM). Each domain was assessed using three separate tests, including letter-, number-, and figure-based material, respectively. Participants completed all tasks on a computer and responded by either typing in words or numbers; using the computer mouse; or pressing keys marked by different colors. Each test included initial practice runs, after which testing followed across several trials. For each of the three tests within a domain, scores were summarized across trials for a measure of overall performance. A composite score of performances across the three tests was calculated and used as the measure of the cognitive domain in question (i.e., episodic memory, working memory). For each of the three tests, scores were summarized across the total number of trials. The three resulting sum scores were z-standardized and averaged to form one composite score for each domain. The standardization has been carried out independently for the training (DyNAMiC) and test (COBRA) samples.

#### 4.2.1. Episodic Memory (EM)

Tests of EM included word recall, number-word recall, and object-location recall. In word recall, participants viewed 16 Swedish concrete nouns, successively appearing on the computer screen. Each word was presented for 6 s, with an inter-stimulus interval (ISI) of 1 s. Directly following encoding, participants reported as many words as they could using the keyboard. Two trials were completed, with a combined maximum score of 32. In number-word recall, participants encoded pairs of 2-digit numbers and concrete plural nouns (e.g., 46 dogs). During encoding, eight number-word pairs were presented, each for 6 s, with an ISI of 1 s. Directly after encoding, nouns were presented again, in re-arranged order, and participants had to report the 2-digit number associated with each presented noun (e.g., How many dogs?). This test included two trials with a total combined maximum score of 16. The third test assessed object-location memory. Participants viewed a 6 × 6 square grid in which 12 objects were consecutively shown at distinct locations. Each object-position pairing was displayed for 8 s, with an ISI of 1 s. Directly following encoding, all objects were simultaneously displayed next to the grid for the participant to place in their correct position within the grid. If unable to recall the correct position of an object, participants had to guess to the best of their ability. Two trials of this test were completed, making the total combined maximum score 24.

### 4.2.2. Working Memory (WM)

Working memory was examined with three tests: letter updating, number updating, and spatial updating. These tests differed from the working memory *n*-back task performed during fMRI scanning. In letter-updating, capital letters (A–D) were consecutively presented on the computer screen, with participants instructed to always keep the three final letters in memory. Each letter was presented for 1 s, with an ISI of 0.5 s. When prompted, at any given moment, participants provided their responses by typing in three letters using the keyboard and provided a guessing-based answer if they failed to remember the correct letters. Four practice trials were followed by 16 test trials consisting of either 7-, 9-, 11-, or 13-letter sequences. The combined maximum number of correct answers across trials was 48 (16 trials × 3 reported letters = 48). The number-updating test followed a columnized 3-back design. Three boxes were presented next to each other on the screen throughout the task, in which a single digit (1–9) was sequentially presented from left to right for 1.5 s with an ISI of 0.5 s. During an ongoing sequence, participants had to judge whether the number currently presented in a specific box matched the last number presented in the same box (i.e., appearing three numbers before). For each presented number, they responded yes/no by pressing assigned keys (“yes” = right index finger; “no” = left index finger). Two practice trials were followed by four test trials, each consisting of 30 numbers. The combined maximum number of correct answers across trials (after discarding responses to the first three numbers in every trial, as these were not preceded by any numbers to be matched with) was 108 (27 numbers × 4 trials). In spatial-updating, participants viewed three 3 × 3 square grids presented next to each other on the computer screen. At the start of each trial, a blue circular object was displayed at a random location within each grid. After 4 s, the circular objects disappeared, leaving the grids empty. An arrow then appeared below each grid, indicating that the circular object in the corresponding grid was to be mentally moved one step in the direction of the arrow. The arrows appeared stepwise from left to right in the grids, each presented for 2.5 s (ISI = 0.5 s). Prompted by three new arrows, participants mentally moved the circular objects one more time, resulting in each circular object having moved two steps from its original location at the end of each trial. Participants indicated which square the circular object in each grid had ended up in using the computer mouse. If unsure, they were instructed to guess. The test was performed across 10 test trials, preceded by five practice trials. The combined maximum number of correct location indications was 30.

### 4.3. Measure of Physical Activity and Cardiovascular Disease Risk

#### 4.3.1. Physical Activity

An extensive battery of self-rating questionnaires was administered in DyNAMiC and COBRA (56, 88). Participants were asked to indicate the frequency (number of hours during a typical summer week; options: 1-14 h with 1-h increments, or 15+ hrs) and the intensity (how physically demanding on a scale from 1 =“not at all” to 5 =“extremely”) by which they typically engage in a selection of activities relevant to life in northern Sweden. These included 15 specific activities. For the present study, we focused on physical activities and on those activities that are purely physical and that individuals are sufficiently engaged in (i.e., physically demanding ≥2.0). Each of these activities was performed by at least 20% of the participants at least once a week; e.g., walking, bicycling, jogging, strength training, household tasks, and daily work-related activities. We computed physical activity frequency (sum hrs/week) to generate physical activity scores accordingly.

#### 4.3.2. Cardiovascular Disease risk

The risk of cardiovascular disease was determined via a multivariable score, according to the algorithm developed in the Framingham Heart Study (122). Variables include age, sex, hypertension diagnosis, systolic blood pressure, body mass index, smoking, and diabetes mellitus. The risk estimates were derived using an algorithm proposed by D’Agostino et al. (122), which employs Cox proportional-hazard regression models to predict the probability of developing any form of cardiovascular disease within 10 years:

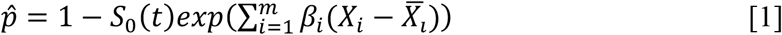

With *S*_*0*_*(t)* being the baseline survival at follow-up *t* (*t* =10 years), *βi* the estimated regression coefficient, X_*i*_ the log-transformed value of the *i*^th^ risk factor, X_*i*_ the corresponding mean, and *m* the number of risk factors included. Baseline survival, means, and regression coefficients were taken from the original algorithm, with DyNAMiC and COBRA participants’ risk variables inserted to compute the final scores. Risk score calculators can be found at *framinghamheartstudy.org*.

### 4.4. Image Acquisitions

Structural, functional, and neurochemical brain measures were acquired using MRI and PET at Umeå University Hospital in northern Sweden. For both DyNAMiC and COBRA, all MRI data were collected using a 3T Discovery MR750 MRI system (General Electric, Healthcare, Illinois, USA) equipped with a 32-channel phased-array head coil. PET was conducted in 3D mode with a Discovery PET/CT 690 (General Electric, WI, United States) to assess whole-brain D1DR with [^11^C]SCH23390 and D2DR with [^11^C]Raclopride at rest in DyNAMiC and COBRA, respectively. Comprehensive descriptions of MRI, PET, and cognitive testing protocols are given elsewhere (56, 88). In this study, we primarily focus on those data directly pertinent to the current investigation.

#### 4.4.1. Functional MRI

For the DyNAMiC dataset, high-resolution anatomical T1-weighted images were collected using a 3-dimensional (3D) fast spoiled gradient-echo sequence with acquisition parameters of 176 sagittal slices, thickness =1 mm, TR =8.2 ms, TE =3.2 ms, flip angle =12°, and a field of view (FOV) =250×250 mm. Whole-brain functional images were acquired during resting state, naturalistic viewing, and an n-back WM task. Functional images were acquired using a T2*-weighted single-shot echo-planar imaging (EPI) sequence, with 330 volumes collected over 12 min. The sequence provided 37 axial slices, slice thickness =3.4 mm, 0.5 mm spacing, TR =2,000 ms, TE =30 ms, flip angle =80°, and FOV =250×250 mm. Ten dummy scans were collected at the start of the sequence. During the resting state, participants were instructed to keep their eyes open and focus on a white fixation cross on a black background displayed on a computer screen through a tilted mirror attached to the head coil. WM was assessed in the scanner (12 min) with a numerical *n*-back task, which consisted of blocks of 1-back, 2-back, and 3-back (102, 103). During movie-watching, the participants viewed and listened to a 12-minute video consisting of selected and chronologically ordered sections from the Swedish movie *Cockpit* (123). Participants were instructed to view the movie attentively and answer a short multiple-choice questionnaire about the movie after the scanning session. We did not monitor participants’ responses to the movie, and the chosen clips were selected to be relatively neutral in emotional content. The storyline follows Valle, a recently fired pilot whose marriage has ended, as he struggles to find new employment. In a desperate attempt to secure a job at an airline specifically recruiting a female pilot, he presents himself as a woman.

In COBRA, data were collected using identical scans for resting state and n-back WM. However, the resting state scan was shorter, lasting only 6 minutes (75, 103).

### 4.5. Functional Connectivity Analysis

Functional data from all conditions (i.e., rest, movie, n-back) were pre-processed using the Statistical Parametric Mapping software package (SPM12). The functional images were first corrected for slice-timing differences and in-scanner motion, followed by registration to anatomical T1 images. Distortion correction was performed using subject-specific and T1 co-registered field maps. The functional time series were subsequently demeaned and detrended, followed by simultaneous nuisance regression and temporal high-pass filtering (threshold at 0.009 Hz) to not re-introduce nuisance signals (124). The nuisance regression model included mean cerebrospinal and white-matter signal and their derivatives, 24-motion parameters (125), a binary vector flagging motion-contaminated volumes exceeding framewise displacement (FD) of 0.2 mm (126), in addition to an 18-parameter RETRICOR model (127, 128) of cardiac pulsation (up to third-order harmonics), respiration (up to fourth-order harmonics), and first-order cardio-respiratory interactions estimated using the PhysIO Toolbox v.5.0 (129). Regression models for n-back included an additional set of finite impulse response (FIR) task regressors (130) to avoid false positive connectivity due to task-evoked activations (131). The FIR regression approach involved fitting mean cross-block responses for each time point within a time-locked window of equal duration to each task block (27 blocks of 20 s), extended by an additional 18 s following each block to account for the duration of the hemodynamic response function (HRF). Given that this approach linearly fits a set of binary task regressors for each time point, it is nearly identical to subtracting the mean task response, with the difference being that FIR regression is better able to handle overlapping task responses and differences in the shape of the HRF (131). The nuisance-regressed images were subsequently normalized to sample-specific group templates (DARTEL) (132) for each dataset, respectively, followed by spatial smoothing using a 6-mm FWHM Gaussian kernel to mitigate DARTEL-induced aliasing and affine-transformed to stereotactic MNI space (ICBM152NLin2009) (133, 134). Functional images from both DyNAMiC and COBRA were preprocessed according to the steps described above, with the only exception that the RETRICOR parameters were excluded from the regression models in COBRA due to technical issues related to the respiration and cardiac traces.

#### 4.5.1. Graph Construction

For all fMRI conditions, functional time series were averaged from 273 cortical and subcortical regions, represented by 5-mm radius spheres, based on a widely employed FC parcellation (89). In addition to 264 regions reported in the Power parcellation, we added 9 additional regions, including some subcortical regions, such as putamen, caudate, and anterior and posterior hippocampus, identified using independent component analysis (8, 135). These regions were categorized into 14 resting state networks according to a consensus partition (89). To mitigate sampling from non-gray matter voxels, each parcel underwent erosion by a permissive gray matter mask (eroding voxels <.1% threshold). The averaged time series were then subjected to Pearson’s correlations, followed by Fisher’s r-to-z transformation, resulting in the creation of a 273×273 adjacency matrix for each participant, with coefficients along the main diagonals set to zero.

To further investigate the impact of network parcellation, we replicated our prediction analysis (**Supplementary Material, Figs S1-S2, and Table S1-S2**) using Schaefer parcellation (92), which entails 300 cortical and subcortical regions.

### 4.6. Deep Neural Network Model

Based on convolutional neural networks, deep learning is an advanced form of artificial intelligence that uses multiple layers of “hidden” neural networks. Deep learning methodologies are capable of automatically identifying complex patterns and representations directly from raw data using these multi-layered networks, thus eliminating the need for explicit feature engineering or manual intervention (136). The success of the training and learning phases depends on the model’s ability to process high-dimensional input data, extracting meaningful features from complex data. This is done while managing the number of trainable parameters, which are crucial for automated feature learning during the construction of the model.

In this study, the inputs to our deep learning models were subject-specific FC maps with a matrix size of 273×273 (e.g., **Fig. 6a**). We generated different versions of each FC map by replacing portions of the network. For example, we relocated the DMN network toward the last nodes, which were assigned as DAN, cerebellar, subcortical, and uncertain in **Figure 6a**. This approach allowed us to create a diverse set of FC matrices for each individual, each reflecting a different composition of edges and neighbors while maintaining the linear relationships exonerated in the original data. Consequently, we augmented the dataset by producing a total of 3,600 FC maps from the initial set of FC maps.

**Figure 6.**
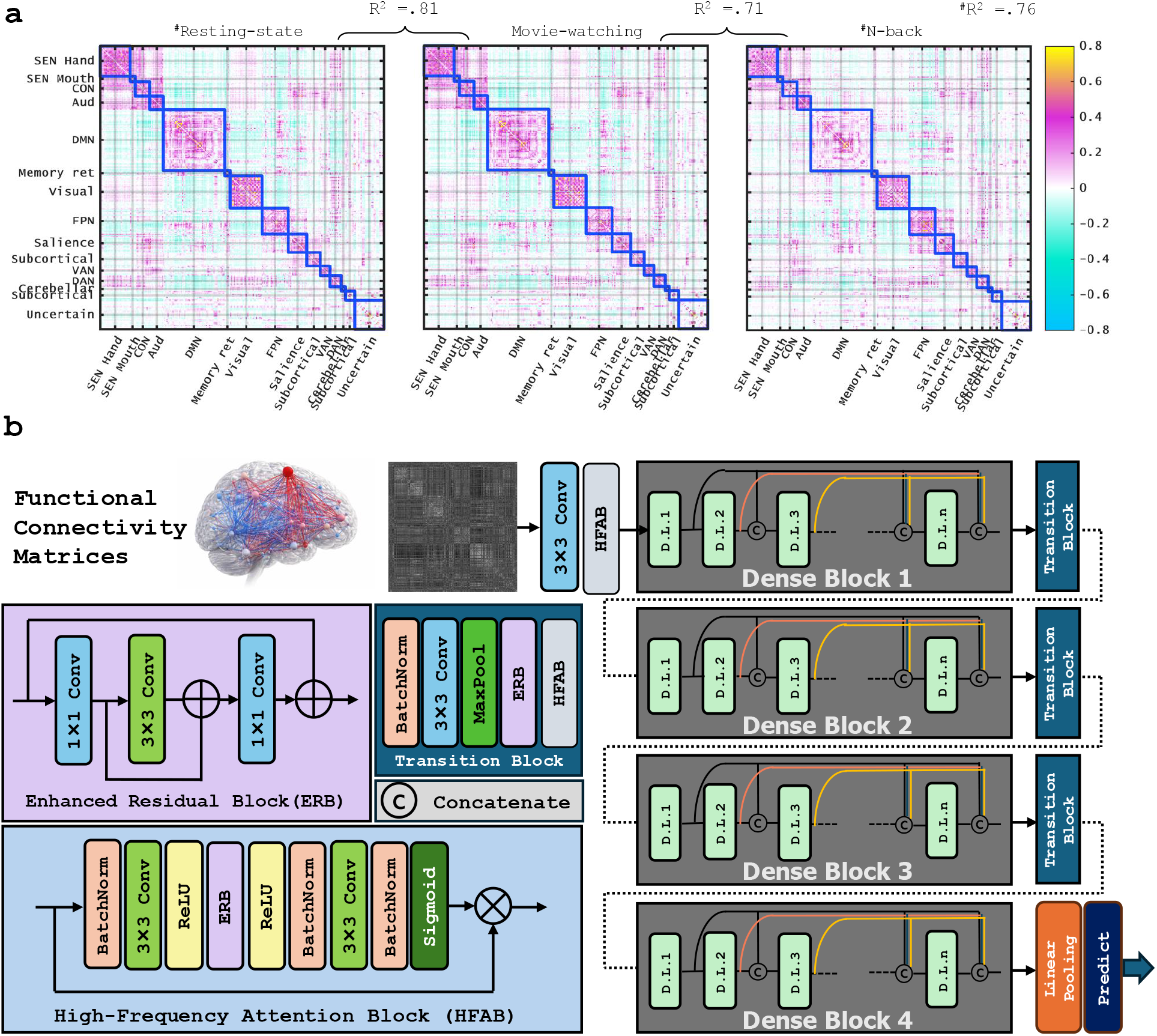
**(a)** Example of a functional connectivity map across three different cognitive states. SEN hand: SENsory hand; SEN Mouth: SENsory Mouth; CON: Cingulo-Operculum control Network; Aud: Auditory; DMN: Default Mode Network; Memory Ret: Memory Retrieval network; Visual: Visual; FPN: Fronto-Parietal Network; Salience: Salience control network; Subcortical (upper row): subcortical network included in original Power parcellation; VAN: Ventral Attention Network; DAN: Dorsal Attention Network; Cerebellar: Cerebellar network; Subcortical (lower row): additional Subcortical regions, including hippocampus and caudate, added to the original Power Parcellation; Uncertain: Regions with less known network assignment. **(b)** DenselyAttention architecture. Enhanced Residual Block (ERB) and High-Frequency Attention Block (HFAB) into the Transition Block. Note that each “D.L.” layer in the table corresponds to the sequence BatchNormalization-ReLU-Conv3×3.

#### Transition

##### Block

Following the DenseNet framework (90), we incorporated the Enhanced Residual Block (ERB) and High-Frequency Attention Block (HFAB) into the dense layers (**Fig. 6b**), termed DenselyAttention, to facilitate feature reuse in each layer. DenseNet (90) diverges from traditional methods like deepening layers or widening network structure by focusing on feature reuse and bypass settings. This results in fewer parameters than similar dense networks such as ResNet (137), enhances feature reuse, improves feature propagation, makes training easier, and reduces issues of gradient vanishing and model degradation.

The architecture of DenseNet is characterized by its dense connectivity pattern, which entails direct connections from each layer to all subsequent layers within its dense block (as illustrated in **Fig. 6b**). This design ensures that every layer has access to the feature maps generated by preceding layers, thereby facilitating a seamless and efficient gradient flow throughout the network. In essence, the knowledge acquired at each layer is propagated forward, enabling the model to effectively capture intricate patterns and dependencies within the data, ultimately enhancing its ability to learn and generalize (90). Additionally, dense blocks provide a regularizing effect, reducing overfitting, particularly on tasks with smaller training datasets (90). This is suitable for this study, which includes relatively small samples. Each sequence combines operations of batch normalization (BN), rectified linear unit (ReLU), and a 3×3 convolution (Conv). Batch normalization can effectively prevent overfitting, as described by equation 2:

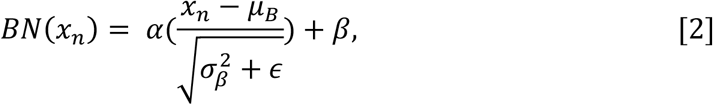

Where *µ*_β_, mean; *σ*_β_, standard deviation; ϵ, random noise; α and β are adaptable variables in training. We utilized the Rectified Linear Unit (ReLU) activation function (138), which activates neurons by directly outputting the input if it is positive, or zero otherwise, as outlined in equation 2.

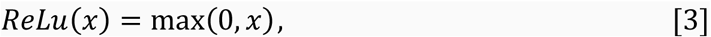

Where *x* is the input to a neuron. ReLU benefits the performance of networks with dense layers by decreasing the computation and selectively optimizing parameters.

Four dense blocks facilitate a stepwise down-sampling in the network. These blocks are connected with transition layers, which consist of a 1×1 convolutional layer followed by a 2×2 average pooling layer.

Additionally, ERB and HFAB pairs are introduced for targeted high-frequency features and residual block enhancement. The ERB-HFAB pairs are stacked sequentially at the beginning of each dense block loop (**Fig. 6b**), with ERB and HFAB having 16 feature maps.

#### High-Frequency Attention Block (HFAB)

Our approach to attentional mechanisms, specifically the HFAB, introduces a sequential attention branch inspired by edge detection. This branch rescales each position based on its neighboring pixels, efficiently focusing on high-frequency areas. The HFAB employs a 3×3 convolution to enhance both the receptive field and computational efficiency. Batch normalization is seamlessly integrated into the attention branch, introducing global statistics during inference without additional computational cost.

#### Enhanced Residual Block (ERB)

We present ERB as an alternative to the traditional residual block. As illustrated in **Figure 6b**, ERB comprises a re-parameterization block (RepBlock) and a ReLU. In the training stage, the RepBlock utilizes a 1×1 convolution to either expand or contract the number of feature maps, employing a 3×3 convolution to extract features in a higher-dimensional space. Furthermore, two skip connections are integrated to mitigate training complexities. During inference, all linear transformations can be unified, facilitating the conversion of each RepBlock into a singular 3×3 convolution. Essentially, ERB capitalizes on the advantages of residual learning.

We implemented model training using Tensorflow 2.11.0 (139) and Keras 2.11.0 (140) as programming interfaces and trained on a fifth-generation MacBook Pro (Apple M1 MAX silicon chips, 10-core CPU, 24-core GPU, 16-core Neural Engine, 64 GB memory). For regression tasks, selecting an appropriate loss function is crucial for guiding the optimization process and ensuring accurate predictions. In this study, we opted for the mean squared error (MSE) loss function due to its suitability for regression problems. To minimize the loss function, we trained the network using the stochastic gradient descent (SGD) optimizer with a learning rate of 8e^-5^ and a Nesterov momentum of 0.9 (141). The number of epochs was set to 100, and the batch size was set to 74. We also added dropout (142) after each convolutional layer, except the first one, with a rate of 0.15.

We conducted training on distinct, identical datasets extracted from DyNAMiC, each comprising FC maps generated from 120 subjects. To prepare each training dataset, we randomly shuffled data for training and validation patches, allocating 80% for training and 20% for validation. For testing, we utilized all FC maps of 60 subjects from the same sample (the age-heterogeneous DyNAMiC study), enabling us to assess model performance through three-fold cross-validation (**Fig. 5**). Each cross-validation fold was a new training with an unseen validation set for the model. Based on its performance on the testing dataset, we selected and employed the winning model for all subsequent analyses. This model, which demonstrated the best performance on the testing dataset, underwent external validation in an independent sample from the age-homogeneous COBRA study.

We initially attempted to predict both episodic memory (EM) and working memory (WM). However, EM prediction was only reliable within and across samples for the resting state, whereas WM prediction generalized most strongly from the movie-watching condition. Because COBRA does not include a movie-watching paradigm, we could not evaluate WM prediction across datasets. For this reason, we focused on EM when examining the brain-cognition.

To explore and visually represent the crucial features of the deep learning models contributing to the prediction of cognitive scores, we used the Grad-CAM (Gradient-weighted Class Activation Mapping) technique (143). This method interprets the model’s decisions by highlighting the regions of the input image with the most significant impact on the model’s output. Grad-CAM conducts a backward pass to compute the gradients of the target class score with respect to the feature maps of the final convolutional layer. We present the average heatmaps calculated for all input data to the model (143). Grad-CAM saliency maps were interpreted qualitatively, with a heuristic threshold (≥ 0.5) applied to highlight regions with relatively higher contribution to the model’s predictions. These values do not reflect statistical significance and should therefore be interpreted descriptively. A further limitation of this study is the absence of ground truth for validating whether highlighted regions truly correspond to the features used by the model during prediction. As such, Grad-CAM provides an approximation of model attention rather than a definitive measure of feature importance. Nevertheless, Grad-CAM remains one of the most widely used and empirically validated interpretability techniques in deep learning, particularly in medical imaging applications. Its integration with established frameworks such as Keras and TensorFlow, together with its ability to generate spatial attributions that align with domain knowledge, makes it a suitable choice for the present study. Future work may incorporate complementary interpretability approaches, including adaptations of the Haufe transformation where applicable to deep learning architectures.

### 4.7. Positron Emission Tomography (PET)

The scanning sessions started with a 5-minute low-dose helical CT scan (20 mA, 120 kV, 0.8 s per revolution), obtained for attenuation correction. During scanning, a thermoplastic mask was attached to the bed surface to minimize head movement.

In DyNAMiC, a 60-minute scan was performed following 350 MBq (337 ± 27 MBq) in list-mode format. Offline re-binning of list-mode data was conducted to achieve a total of 49 frames with increasing length. In COBRA, a 55-min, 18-frame dynamic PET scan was acquired during rest following intravenous bolus injection of approximately 250 MBq 11C-raclopride (264 ± 19 MBq). For both studies, attenuation- and decay-corrected images (47 slices, field of view = 25 cm, 256×256-pixel transaxial images, voxel size = 0.977× 0.97×3.27 mm^3^) were reconstructed with the iterative VUE Point HD-SharpIR algorithm (GE; 6 iterations, 24 subsets, 3.0 mm post filtering; full-width-at-half-maximum (FWHM): 3.2 mm). The estimation of receptor availability or binding potential relative to non-displaceable binding (BP_ND_) was carried out following previously described procedures with the cerebellum as a reference region (76). PET images were motion-corrected and co-registered with the structural T1-weighted images from the corresponding session using Statistical Parametric Mapping software (SPM12, Wellcome Department of Imaging Science, Functional Imaging Laboratory, London, UK). Motion-corrected PET data were resliced to match the spatial dimensions of MR data (1 mm^3^ isotropic, 256×256×256). The mean of the first five frames was used as a source for co-registration. In DyNAMiC, frame-to-frame head motion correction, with translations ranging from 0.23 to 4.22 mm (mean ± sd =0.95 ± 0.54 mm), revealed a trend-level difference across age-groups (age <40 and age ≥40 years), as determined by Student’s t-test (*t* =2.0, *p* =.047; mean ± sd for younger individuals = 1.07 ± 0.52, mean ± sd for older individuals =0.90 ± 0.55). Partial-volume-effect (PVE) correction was achieved using the symmetric geometric transfer matrix (SGTM) method for regional correction, implemented in FreeSurfer (144). An estimated point-spread-function of 2.5 mm FWHM was utilized. Regional estimates of BP_ND_ were calculated within the automated FreeSurfer segmentations employing the simplified reference tissue model (SRTM (145)). In the current study, we focused on the striatal BP_ND_, calculated as an average of BP_ND_ across the caudate and putamen.

### 4.8. The direct impact of dopamine on BCG through mediation analysis

To evaluate whether functional variability mediates the relationship between D1DR and prediction gap connectivity, we conducted additional mediation analyses. We first computed entropy, which estimates within-region signal variability (86), and then averaged this measure across all striatal regions (left caudate: MNI coordinate ≈ [–12, 12, 6], right caudate: MNI coordinate ≈ [10, 14, 2], left putamen: MNI coordinates ≈ [–24, 6, 4], right putamen: MNI coordinates ≈ [29, 1, 4].). Following the mediation analysis framework proposed by Baron and Kenny (146), our goal was to determine whether the association between D1/D2 receptors and BCG is mediated by regional variability (entropy) or if the indirect effect exceeds the direct association between D1/D2 receptors and BCG. To assess the statistical significance of this mediation effect, we employed the bootstrapping method as outlined by Preacher and Hayes (147), and age has been controlled for in all statistical analyses.

### 4.9. Statistical significance analysis

Statistical analyses were carried out using SPSS (IBM Corp., V24.0.0, Armonk, NY, USA), MATLAB (The MathWorks Inc., V9.13.0 (R2022b), Natick, MA, USA), and GraphPad Prism (GraphPad Software, Inc., V5.01, CA, USA). We performed partial correlations between predicted and actual scores, as well as linear regression analyses. To investigate the relationship between generated gap variables and DA receptor availability, we controlled for age (in DyNAMiC) using partial correlation. The Mann-Whitney *U* test was used to calculate the mean differences in prediction accuracy. The level of statistical significance was set at *p-value* ≤0.05. For the bootstrap-based comparison of model performance (bootstrap resampling with 1000 iterations), no test statistic with an associated degree of freedom is reported. Instead, statistical inference is based on the bootstrap distribution of the difference in correlation coefficients (Δr) and its 95% confidence interval. As bootstrap confidence-interval-based inference does not rely on an analytic sampling distribution, degrees of freedom are not defined for this procedure.

Out-of-sample predictive performance was quantified using the coefficient of determination (*r*^2^) computed via a sum-of-squares formulation (148). Unlike squared correlation coefficients, which capture only linear association, this metric evaluates how well model predictions approximate observed values relative to a baseline model. Specifically, out-of-sample *r*^2^ was defined as

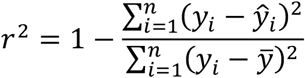

where *y*_i_ denotes the observed outcome in the test set, 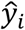 the corresponding model prediction, and 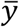 train denotes the mean of the outcome variable in the training set. Using the training-set mean as the baseline ensures a strictly out-of-sample evaluation and avoids information leakage. Under this formulation, positive *r*^2^ values indicate performance exceeding the null model (predicting the training mean), whereas negative values indicate worse-than-baseline performance. Because this formulation directly compares prediction error to baseline variance, it provides a more appropriate measure of predictive accuracy than correlation-based metrics, particularly in the presence of scale or offset differences between predicted and observed values (148).

## Acknowledgments

This work was funded by the Swedish Research Council (grant number 2021-02558), Knut and Alice Wallenberg Foundation (Wallenberg Fellow grant to A.S.), Bank of Sweden (RJ, grant number P20-0515 to A.S.), StratNeuro grant at Karolinska Institute (A.S.). Morteza Esmaeili and Erin Bjørkeli were supported by the Southern Eastern Norway Regional Health Authority (Helse Sør-øst RHF, HSø, grant numbers 2018047 and 2021023, respectively).

## REFERENCES

1. Biswal B, Yetkin FZ, Haughton VM, Hyde JS. Functional connectivity in the motor cortex of resting human brain using echo-planar MRI. Magn Reson Med. 1995;34(4):537–41.

2. Buckner RL, Krienen FM. The evolution of distributed association networks in the human brain. Trends Cogn Sci. 2013;17(12):648–65.

3. van den Heuvel MP, Hulshoff Pol HE. Exploring the brain network: a review on resting-state fMRI functional connectivity. European neuropsychopharmacology : the journal of the European College of Neuropsychopharmacology. 2010;20(8):519–34.

4. Ferreira LK, Busatto GF. Resting-state functional connectivity in normal brain aging. Neurosci Biobehav Rev. 2013;37(3):384–400.

5. Damoiseaux JS. Effects of aging on functional and structural brain connectivity. Neuroimage. 2017;160:32–40.

6. Fox MD, Greicius M. Clinical applications of resting state functional connectivity. Front Syst Neurosci. 2010;4:19.

7. Andrews-Hanna JR, Snyder AZ, Vincent JL, Lustig C, Head D, Raichle ME, et al. Disruption of large-scale brain systems in advanced aging. Neuron. 2007;56(5):924–35.

8. Salami A, Wahlin A, Kaboodvand N, Lundquist A, Nyberg L. Longitudinal Evidence for Dissociation of Anterior and Posterior MTL Resting-State Connectivity in Aging: Links to Perfusion and Memory. Cereb Cortex. 2016;26(10):3953–63.

9. Kaboodvand N, Backman L, Nyberg L, Salami A. The retrosplenial cortex: A memory gateway between the cortical default mode network and the medial temporal lobe. Hum Brain Mapp. 2018;39(5):2020–34.

10. Seeley WW, Menon V, Schatzberg AF, Keller J, Glover GH, Kenna H, et al. Dissociable intrinsic connectivity networks for salience processing and executive control. The Journal of neuroscience : the official journal of the Society for Neuroscience. 2007;27(9):2349–56.

11. Avelar-Pereira B, Backman L, Wahlin A, Nyberg L, Salami A. Age-Related Differences in Dynamic Interactions Among Default Mode, Frontoparietal Control, and Dorsal Attention Networks during Resting-State and Interference Resolution. Front Aging Neurosci. 2017;9:152.

12. Avelar-Pereira B, Backman L, Wahlin A, Nyberg L, Salami A. Increased functional homotopy of the prefrontal cortex is associated with corpus callosum degeneration and working memory decline. Neurobiol Aging. 2020;96:68–78.

13. Machner B, Braun L, Imholz J, Koch PJ, Munte TF, Helmchen C, et al. Resting-State Functional Connectivity in the Dorsal Attention Network Relates to Behavioral Performance in Spatial Attention Tasks and May Show Task-Related Adaptation. Front Hum Neurosci. 2021;15:757128.

14. Schimmelpfennig J, Topczewski J, Zajkowski W, Jankowiak-Siuda K. The role of the salience network in cognitive and affective deficits. Front Hum Neurosci. 2023;17:1133367.

15. Ferreira LK, Regina AC, Kovacevic N, Martin Mda G, Santos PP, Carneiro Cde G, et al. Aging Effects on Whole-Brain Functional Connectivity in Adults Free of Cognitive and Psychiatric Disorders. Cereb Cortex. 2016;26(9):3851–65.

16. Smith SM, Elliott LT, Alfaro-Almagro F, McCarthy P, Nichols TE, Douaud G, et al. Brain aging comprises many modes of structural and functional change with distinct genetic and biophysical associations. Elife. 2020;9.

17. Shen X, Finn ES, Scheinost D, Rosenberg MD, Chun MM, Papademetris X, et al. Using connectome-based predictive modeling to predict individual behavior from brain connectivity. Nat Protoc. 2017;12(3):506–18.

18. Vul E, Harris C, Winkielman P, Pashler H. Puzzlingly High Correlations in fMRI Studies of Emotion, Personality, and Social Cognition. Perspect Psychol Sci. 2009;4(3):274–90.

19. Bzdok D, Eickenberg M, Varoquaux G, Thirion B. Hierarchical Region-Network Sparsity for High-Dimensional Inference in Brain Imaging. Inf Process Med Imaging. 2017;10265:323–35.

20. Bzdok D, Meyer-Lindenberg A. Machine Learning for Precision Psychiatry: Opportunities and Challenges. Biol Psychiatry Cogn Neurosci Neuroimaging. 2018;3(3):223–30.

21. Bzdok D, Krzywinski M, Altman N. Machine learning: supervised methods. Nat Methods. 2018;15(1):5–6.

22. Sui J, Jiang R, Bustillo J, Calhoun V. Neuroimaging-based Individualized Prediction of Cognition and Behavior for Mental Disorders and Health: Methods and Promises. Biol Psychiatry. 2020;88(11):818–28.

23. Vieira S, Liang X, Guiomar R, Mechelli A. Can we predict who will benefit from cognitive-behavioural therapy? A systematic review and meta-analysis of machine learning studies. Clin Psychol Rev. 2022;97:102193.

24. Plis SM, Hjelm DR, Salakhutdinov R, Allen EA, Bockholt HJ, Long JD, et al. Deep learning for neuroimaging: a validation study. Front Neurosci. 2014;8:229.

25. van der Burgh HK, Schmidt R, Westeneng HJ, de Reus MA, van den Berg LH, van den Heuvel MP. Deep learning predictions of survival based on MRI in amyotrophic lateral sclerosis. Neuroimage Clin. 2017;13:361–9.

26. Nguyen H, Nguyen V, Nguyen T, Larsen ME, O’Dea B, Nguyen DT, et al., editors. Jointly Predicting Affective and Mental Health Scores Using Deep Neural Networks of Visual Cues on the Web2018; Cham: Springer International Publishing.

27. Li H, Jiang G, Zhang J, Wang R, Wang Z, Zheng WS, et al. Fully convolutional network ensembles for white matter hyperintensities segmentation in MR images. Neuroimage. 2018;183:650–65.

28. Choi Y, Kwon Y, Lee H, Kim BJ, Paik MC, Won J-H, editors. Ensemble of Deep Convolutional Neural Networks for Prognosis of Ischemic Stroke. Brainlesion: Glioma, Multiple Sclerosis, Stroke and Traumatic Brain Injuries; 2016 2016//; Cham: Springer International Publishing.

29. He T, Kong R, Holmes AJ, Nguyen M, Sabuncu MR, Eickhoff SB, et al. Deep neural networks and kernel regression achieve comparable accuracies for functional connectivity prediction of behavior and demographics. Neuroimage. 2020;206:116276.

30. Vieira BH, Schöttner M, Calhoun VD, Salmon CEG. Beyond functional connectivity: deep learning applied to resting-state fMRI time series in the prediction of 58 human traits in the HCP. 2024.

31. Kong J, Wolcott E, Wang Z, Jorgenson K, Harvey WF, Tao J, et al. Altered resting state functional connectivity of the cognitive control network in fibromyalgia and the modulation effect of mind-body intervention. Brain Imaging Behav. 2019;13(2):482–92.

32. Sripada C, Angstadt M, Taxali A, Clark DA, Greathouse T, Rutherford S, et al. Brain-wide functional connectivity patterns support general cognitive ability and mediate effects of socioeconomic status in youth. Transl Psychiatry. 2021;11(1):571.

33. Chen J, Tam A, Kebets V, Orban C, Ooi LQR, Asplund CL, et al. Shared and unique brain network features predict cognitive, personality, and mental health scores in the ABCD study. Nat Commun. 2022;13(1):2217.

34. Omidvarnia A, Sasse L, Larabi DI, Raimondo F, Hoffstaedter F, Kasper J, et al. Is resting state fMRI better than individual characteristics at predicting cognition? bioRxiv. 2023:2023.02.18.529076.

35. Jiang L, Qiao K, Li C. Distance-based functional criticality in the human brain: intelligence and emotional intelligence. BMC Bioinformatics. 2021;22(1):32.

36. Vieira BH, Pamplona GSP, Fachinello K, Silva AK, Foss MP, Salmon CEG. On the prediction of human intelligence from neuroimaging: A systematic review of methods and reporting. Intelligence. 2022;93:101654.

37. Jangraw DC, Gonzalez-Castillo J, Handwerker DA, Ghane M, Rosenberg MD, Panwar P, et al. A functional connectivity-based neuromarker of sustained attention generalizes to predict recall in a reading task. Neuroimage. 2018;166:99–109.

38. Chen Q, Beaty RE, Cui Z, Sun J, He H, Zhuang K, et al. Brain hemispheric involvement in visuospatial and verbal divergent thinking. Neuroimage. 2019;202:116065.

39. Rosenberg MD, Finn ES, Scheinost D, Papademetris X, Shen X, Constable RT, et al. A neuromarker of sustained attention from whole-brain functional connectivity. Nat Neurosci. 2016;19(1):165–71.

40. Beaty RE, Kenett YN, Christensen AP, Rosenberg MD, Benedek M, Chen Q, et al. Robust prediction of individual creative ability from brain functional connectivity. Proc Natl Acad Sci U S A. 2018;115(5):1087–92.

41. Hsu WT, Rosenberg MD, Scheinost D, Constable RT, Chun MM. Resting-state functional connectivity predicts neuroticism and extraversion in novel individuals. Soc Cogn Affect Neurosci. 2018;13(2):224–32.

42. Gabrieli JDE, Ghosh SS, Whitfield-Gabrieli S. Prediction as a humanitarian and pragmatic contribution from human cognitive neuroscience. Neuron. 2015;85(1):11–26.

43. Finn ES, Todd Constable R. Individual variation in functional brain connectivity: implications for personalized approaches to psychiatric disease. Dialogues Clin Neurosci. 2016;18(3):277–87.

44. Woo CW, Chang LJ, Lindquist MA, Wager TD. Building better biomarkers: brain models in translational neuroimaging. Nat Neurosci. 2017;20(3):365–77.

45. Finn ES, Shen X, Scheinost D, Rosenberg MD, Huang J, Chun MM, et al. Functional connectome fingerprinting: identifying individuals using patterns of brain connectivity. Nat Neurosci. 2015;18(11):1664–71.

46. Finn ES, Scheinost D, Finn DM, Shen X, Papademetris X, Constable RT. Can brain state be manipulated to emphasize individual differences in functional connectivity? Neuroimage. 2017;160:140–51.

47. Grady CL, Ryan JD. Age-Related Differences in the Human Hippocampus: Behavioral, Structural and Functional Measures. In: Hannula DE, Duff MC, editors. The Hippocampus from Cells to Systems: Structure, Connectivity, and Functional Contributions to Memory and Flexible Cognition. Cham: Springer International Publishing; 2017. p. 167–208.

48. Campbell KL, Schacter DL. Aging and the Resting State: Cognition is not Obsolete. Lang Cogn Neurosci. 2017;32(6):692–4.

49. Geerligs L, Tsvetanov KA. The use of resting state data in an integrative approach to studying neurocognitive ageing - Commentary on Campbell and Schacter (2016). Lang Cogn Neurosci. 2017;32(6):684–91.

50. Tavor I, Parker Jones O, Mars RB, Smith SM, Behrens TE, Jbabdi S. Task-free MRI predicts individual differences in brain activity during task performance. Science. 2016;352(6282):216–20.

51. Greene AS, Gao S, Scheinost D, Constable RT. Task-induced brain state manipulation improves prediction of individual traits. Nat Commun. 2018;9(1):2807.

52. Finn ES, Bandettini PA. Movie-watching outperforms rest for functional connectivity-based prediction of behavior. Neuroimage. 2021;235:117963.

53. Finn ES, Glerean E, Hasson U, Vanderwal T. Naturalistic imaging: The use of ecologically valid conditions to study brain function. Neuroimage. 2022;247:118776.

54. Hasson U, Nir Y, Levy I, Fuhrmann G, Malach R. Intersubject synchronization of cortical activity during natural vision. Science. 2004;303(5664):1634–40.

55. Geerligs L, Rubinov M, Cam C, Henson RN. State and Trait Components of Functional Connectivity: Individual Differences Vary with Mental State. The Journal of neuroscience : the official journal of the Society for Neuroscience. 2015;35(41):13949–61.

56. Nordin K, Gorbach T, Pedersen R, Panes Lundmark V, Johansson J, Andersson M, et al. DyNAMiC: A prospective longitudinal study of dopamine and brain connectomes: A new window into cognitive aging. J Neurosci Res. 2022;100(6):1296–320.

57. Cattell RB. Abilities: Their structure, growth, and action. 1971.

58. Kyllonen PC. Is working memory capacity Spearman’s g? Human abilities: Psychology Press; 2013. p. 49–75.

59. Unsworth N, Fukuda K, Awh E, Vogel EK. Working memory and fluid intelligence: capacity, attention control, and secondary memory retrieval. Cogn Psychol. 2014;71:1–26.

60. Tulving E, Schacter DL. Priming and human memory systems. Science. 1990;247(4940):301–6.

61. Smallwood J, Schooler JW. The science of mind wandering: empirically navigating the stream of consciousness. Annu Rev Psychol. 2015;66:487–518.

62. Dosenbach NU, Nardos B, Cohen AL, Fair DA, Power JD, Church JA, et al. Prediction of individual brain maturity using fMRI. Science. 2010;329(5997):1358–61.

63. Cole JH, Franke K. Predicting Age Using Neuroimaging: Innovative Brain Ageing Biomarkers. Trends Neurosci. 2017;40(12):681–90.

64. Dunas T, Wahlin A, Nyberg L, Boraxbekk CJ. Multimodal Image Analysis of Apparent Brain Age Identifies Physical Fitness as Predictor of Brain Maintenance. Cereb Cortex. 2021;31(7):3393–407.

65. Farnsworth von Cederwald B, Josefsson M, Wahlin A, Nyberg L, Karalija N. Association of Cardiovascular Risk Trajectory With Cognitive Decline and Incident Dementia. Neurology. 2022;98(20):e2013–e22.

66. Jonasson LS, Nyberg L, Kramer AF, Lundquist A, Riklund K, Boraxbekk CJ. Aerobic Exercise Intervention, Cognitive Performance, and Brain Structure: Results from the Physical Influences on Brain in Aging (PHIBRA) Study. Front Aging Neurosci. 2016;8:336.

67. Tetereva A, Pat N. Brain age has limited utility as a biomarker for capturing fluid cognition in older individuals. eLife. 2024;12:RP87297.

68. Nyberg L, Lovden M, Riklund K, Lindenberger U, Backman L. Memory aging and brain maintenance. Trends Cogn Sci. 2012;16(5):292–305.

69. Karalija N, Wahlin A, Ek J, Rieckmann A, Papenberg G, Salami A, et al. Cardiovascular factors are related to dopamine integrity and cognition in aging. Ann Clin Transl Neurol. 2019;6(11):2291–303.

70. Backman L, Lindenberger U, Li SC, Nyberg L. Linking cognitive aging to alterations in dopamine neurotransmitter functioning: recent data and future avenues. Neurosci Biobehav Rev. 2010;34(5):670–7.

71. Bäckman L, Nyberg L, Soveri A, Johansson J, Andersson M, Dahlin E, et al. Effects of working-memory training on striatal dopamine release. Science. 2011;333(6043):718.

72. Volkow ND, Gur RC, Wang GJ, Fowler JS, Moberg PJ, Ding YS, et al. Association between decline in brain dopamine activity with age and cognitive and motor impairment in healthy individuals. Am J Psychiatry. 1998;155(3):344–9.

73. Juarez EJ, Samanez-Larkin GR. Exercise, Dopamine, and Cognition in Older Age. Trends Cogn Sci. 2019;23(12):986–8.

74. de Boer L, Axelsson J, Riklund K, Nyberg L, Dayan P, Backman L, et al. Attenuation of dopamine-modulated prefrontal value signals underlies probabilistic reward learning deficits in old age. Elife. 2017;6.

75. Nyberg L, Karalija N, Salami A, Andersson M, Wahlin A, Kaboovand N, et al. Dopamine D2 receptor availability is linked to hippocampal-caudate functional connectivity and episodic memory. Proc Natl Acad Sci U S A. 2016;113(28):7918–23.

76. Nordin K, Nyberg L, Andersson M, Karalija N, Riklund K, Backman L, et al. Distinct and Common Large-Scale Networks of the Hippocampal Long Axis in Older Age: Links to Episodic Memory and Dopamine D2 Receptor Availability. Cereb Cortex. 2021;31(7):3435–50.

77. Papenberg G, Karalija N, Salami A, Rieckmann A, Andersson M, Axelsson J, et al. Balance between Transmitter Availability and Dopamine D2 Receptors in Prefrontal Cortex Influences Memory Functioning. Cereb Cortex. 2020;30(3):989–1000.

78. Cools R, D’Esposito M. Inverted-U-shaped dopamine actions on human working memory and cognitive control. Biol Psychiatry. 2011;69(12):e113–25.

79. Zahrt J, Taylor JR, Mathew RG, Arnsten AF. Supranormal stimulation of D1 dopamine receptors in the rodent prefrontal cortex impairs spatial working memory performance. The Journal of neuroscience : the official journal of the Society for Neuroscience. 1997;17(21):8528–35.

80. Karalija N, Papenberg G, Johansson J, Wahlin A, Salami A, Andersson M, et al. Longitudinal support for the correlative triad among aging, dopamine D2-like receptor loss, and memory decline. Neurobiol Aging. 2024;136:125–32.

81. Servan-Schreiber D, Printz H, Cohen JD. A network model of catecholamine effects: gain, signal-to-noise ratio, and behavior. Science. 1990;249(4971):892–5.

82. Seamans JK, Yang CR. The principal features and mechanisms of dopamine modulation in the prefrontal cortex. Prog Neurobiol. 2004;74(1):1–58.

83. Guitart-Masip M, Salami A, Garrett D, Rieckmann A, Lindenberger U, Backman L. BOLD Variability is Related to Dopaminergic Neurotransmission and Cognitive Aging. Cereb Cortex. 2016;26(5):2074–83.

84. Li SC, Rieckmann A. Neuromodulation and aging: implications of aging neuronal gain control on cognition. Curr Opin Neurobiol. 2014;29:148–58.

85. Pedersen R, Johansson J, Salami A. Dopamine D1-signaling modulates maintenance of functional network segregation in aging. Aging Brain. 2023;3:100079.

86. Shafiei G, Zeighami Y, Clark CA, Coull JT, Nagano-Saito A, Leyton M, et al. Dopamine Signaling Modulates the Stability and Integration of Intrinsic Brain Networks. Cereb Cortex. 2019;29(1):397–409.

87. Fan L, Su J, Qin J, Hu D, Shen H. A Deep Network Model on Dynamic Functional Connectivity With Applications to Gender Classification and Intelligence Prediction. Front Neurosci. 2020;14:881.

88. Nevalainen N, Riklund K, Andersson M, Axelsson J, Ogren M, Lovden M, et al. COBRA: A prospective multimodal imaging study of dopamine, brain structure and function, and cognition. Brain Res. 2015;1612:83– 103.

89. Power JD, Cohen AL, Nelson SM, Wig GS, Barnes KA, Church JA, et al. Functional network organization of the human brain. Neuron. 2011;72(4):665–78.

90. Huang G, Liu Z, van der Maaten L, Weinberger KQ. Densely Connected Convolutional Networks. 160806993. 2016.

91. Zhao W, Makowski C, Hagler DJ, Garavan HP, Thompson WK, Greene DJ, et al. Task fMRI paradigms may capture more behaviorally relevant information than resting-state functional connectivity. Neuroimage. 2023;270:119946.

92. Schaefer A, Kong R, Gordon EM, Laumann TO, Zuo XN, Holmes AJ, et al. Local-Global Parcellation of the Human Cerebral Cortex from Intrinsic Functional Connectivity MRI. Cereb Cortex. 2018;28(9):3095–114.

93. Kohncke Y, Papenberg G, Jonasson L, Karalija N, Wahlin A, Salami A, et al. Self-rated intensity of habitual physical activities is positively associated with dopamine D(2/3) receptor availability and cognition. Neuroimage. 2018;181:605–16.

94. Kurkela K, Ritchey M. Intrinsic functional connectivity among memory networks does not predict individual differences in narrative recall. Imaging Neuroscience. 2024;2:1–17.

95. Finn ES. Is it time to put rest to rest? Trends in Cognitive Sciences. 2021;25(12):1021–32.

96. Zhu Y, Zang F, Wang Q, Zhang Q, Tan C, Zhang S, et al. Connectome-based model predicts episodic memory performance in individuals with subjective cognitive decline and amnestic mild cognitive impairment. Behav Brain Res. 2021;411:113387.

97. Wahlheim CN, Christensen AP, Reagh ZM, Cassidy BS. Intrinsic functional connectivity in the default mode network predicts mnemonic discrimination: A connectome-based modeling approach. Hippocampus. 2022;32(1):21–37.

98. Gonzalez-Castillo J, Kam JWY, Hoy CW, Bandettini PA. How to Interpret Resting-State fMRI: Ask Your Participants. The Journal of neuroscience : the official journal of the Society for Neuroscience. 2021;41(6):1130– 41.

99. Stawarczyk D, D’Argembeau A. Neural correlates of personal goal processing during episodic future thinking and mind-wandering: An ALE meta-analysis. Hum Brain Mapp. 2015;36(8):2928–47.

100. Blonde P, Sperduti M, Makowski D, Piolino P. Bored, distracted, and forgetful: The impact of mind wandering and boredom on memory encoding. Q J Exp Psychol (Hove). 2022;75(1):53–69.

101. Karapanagiotidis T, Bernhardt BC, Jefferies E, Smallwood J. Tracking thoughts: Exploring the neural architecture of mental time travel during mind-wandering. Neuroimage. 2017;147:272–81.

102. Salami A, Garrett DD, Wahlin A, Rieckmann A, Papenberg G, Karalija N, et al. Dopamine D(2/3) Binding Potential Modulates Neural Signatures of Working Memory in a Load-Dependent Fashion. The Journal of neuroscience : the official journal of the Society for Neuroscience. 2019;39(3):537–47.

103. Salami A, Rieckmann A, Karalija N, Avelar-Pereira B, Andersson M, Wahlin A, et al. Neurocognitive Profiles of Older Adults with Working-Memory Dysfunction. Cereb Cortex. 2018;28(7):2525–39.

104. Eriksson J, Vogel EK, Lansner A, Bergstrom F, Nyberg L. Neurocognitive Architecture of Working Memory. Neuron. 2015;88(1):33–46.

105. Liang X, Zou Q, He Y, Yang Y. Topologically Reorganized Connectivity Architecture of Default-Mode, Executive-Control, and Salience Networks across Working Memory Task Loads. Cereb Cortex. 2016;26(4):1501– 11.

106. Gilson M, Deco G, Friston KJ, Hagmann P, Mantini D, Betti V, et al. Effective connectivity inferred from fMRI transition dynamics during movie viewing points to a balanced reconfiguration of cortical interactions. Neuroimage. 2018;180(Pt B):534–46.

107. Betzel RF, Byrge L, Esfahlani FZ, Kennedy DP. Temporal fluctuations in the brain’s modular architecture during movie-watching. Neuroimage. 2020;213:116687.

108. Zhang H, Zhao R, Hu X, Guan S, Margulies DS, Meng C, et al. Cortical connectivity gradients and local timescales during cognitive states are modulated by cognitive loads. Brain Struct Funct. 2022;227(8):2701–12.

109. Ito T, Brincat SL, Siegel M, Mill RD, He BJ, Miller EK, et al. Task-evoked activity quenches neural correlations and variability across cortical areas. PLoS Comput Biol. 2020;16(8):e1007983.

110. Tian YE, Cropley V, Maier AB, Lautenschlager NT, Breakspear M, Zalesky A. Heterogeneous aging across multiple organ systems and prediction of chronic disease and mortality. Nat Med. 2023;29(5):1221–31.

111. Arida RM, Teixeira-Machado L. The Contribution of Physical Exercise to Brain Resilience. Front Behav Neurosci. 2020;14:626769.

112. Klil-Drori S, Cinalioglu K, Rej S. Brain Health and the Role of Exercise in Maintaining Late-Life Cognitive Reserve: A Narrative Review Providing the Neuroprotective Mechanisms of Exercise. The American Journal of Geriatric Psychiatry. 2022;30(4, Supplement):S72.

113. Nithianantharajah J, Hannan AJ. The neurobiology of brain and cognitive reserve: Mental and physical activity as modulators of brain disorders. Progress in Neurobiology. 2009;89(4):369–82.

114. Cabeza R, Albert M, Belleville S, Craik FIM, Duarte A, Grady CL, et al. Maintenance, reserve and compensation: the cognitive neuroscience of healthy ageing. Nat Rev Neurosci. 2018;19(11):701–10.

115. Korkki SM, Johansson J, Nordin K, Pedersen R, Bäckman L, Rieckmann A, et al. Dedifferentiation of caudate functional organization is linked to reduced D1 dopamine receptor availability and poorer memory function in aging. bioRxiv. 2024:2024.03.18.585623.

116. Berry AS, Shah VD, Furman DJ, White RL, 3rd, Baker SL, O’Neil JP, et al. Dopamine Synthesis Capacity is Associated with D2/3 Receptor Binding but Not Dopamine Release. Neuropsychopharmacology. 2018;43(6):1201–11.

117. Volkow ND, Wang GJ, Fowler JS, Ding YS, Gur RC, Gatley J, et al. Parallel loss of presynaptic and postsynaptic dopamine markers in normal aging. Ann Neurol. 1998;44(1):143–7.

118. Kienast T, Siessmeier T, Wrase J, Braus DF, Smolka MN, Buchholz HG, et al. Ratio of dopamine synthesis capacity to D2 receptor availability in ventral striatum correlates with central processing of affective stimuli. Eur J Nucl Med Mol Imaging. 2008;35(6):1147–58.

119. Heinz A, Siessmeier T, Wrase J, Buchholz HG, Grunder G, Kumakura Y, et al. Correlation of alcohol craving with striatal dopamine synthesis capacity and D2/3 receptor availability: a combined [18F]DOPA and [18F]DMFP PET study in detoxified alcoholic patients. Am J Psychiatry. 2005;162(8):1515–20.

120. Ito H, Kodaka F, Takahashi H, Takano H, Arakawa R, Shimada H, et al. Relation between presynaptic and postsynaptic dopaminergic functions measured by positron emission tomography: implication of dopaminergic tone. The Journal of neuroscience : the official journal of the Society for Neuroscience. 2011;31(21):7886–90.

121. Nyberg L, Andersson M, Lundquist A, Baare WFC, Bartres-Faz D, Bertram L, et al. Individual differences in brain aging: heterogeneity in cortico-hippocampal but not caudate atrophy rates. Cereb Cortex. 2023;33(9):5075– 81.

122. D’Agostino RB, Sr., Vasan RS, Pencina MJ, Wolf PA, Cobain M, Massaro JM, et al. General cardiovascular risk profile for use in primary care: the Framingham Heart Study. Circulation. 2008;117(6):743–53.

123. Johansson J, Nordin K, Pedersen R, Karalija N, Papenberg G, Andersson M, et al. Biphasic patterns of age-related differences in dopamine D1 receptors across the adult lifespan. Cell Rep. 2023;42(9):113107.

124. Hallquist MN, Hwang K, Luna B. The nuisance of nuisance regression: spectral misspecification in a common approach to resting-state fMRI preprocessing reintroduces noise and obscures functional connectivity. Neuroimage. 2013;82:208–25.

125. Friston KJ, Williams S, Howard R, Frackowiak RS, Turner R. Movement-related effects in fMRI time-series. Magn Reson Med. 1996;35(3):346–55.

126. Power JD, Barnes KA, Snyder AZ, Schlaggar BL, Petersen SE. Spurious but systematic correlations in functional connectivity MRI networks arise from subject motion. Neuroimage. 2012;59(3):2142–54.

127. Glover GH, Li TQ, Ress D. Image-based method for retrospective correction of physiological motion effects in fMRI: RETROICOR. Magn Reson Med. 2000;44(1):162–7.

128. Hutton C, Josephs O, Stadler J, Featherstone E, Reid A, Speck O, et al. The impact of physiological noise correction on fMRI at 7 T. Neuroimage. 2011;57(1):101–12.

129. Kasper L, Bollmann S, Diaconescu AO, Hutton C, Heinzle J, Iglesias S, et al. The PhysIO Toolbox for Modeling Physiological Noise in fMRI Data. J Neurosci Methods. 2017;276:56–72.

130. Fair DA, Schlaggar BL, Cohen AL, Miezin FM, Dosenbach NU, Wenger KK, et al. A method for using blocked and event-related fMRI data to study “resting state” functional connectivity. Neuroimage. 2007;35(1):396–405.

131. Cole MW, Ito T, Schultz D, Mill R, Chen R, Cocuzza C. Task activations produce spurious but systematic inflation of task functional connectivity estimates. Neuroimage. 2019;189:1–18.

132. Ashburner J. A fast diffeomorphic image registration algorithm. Neuroimage. 2007;38(1):95–113.

133. Fonov V, Evans AC, Botteron K, Almli CR, McKinstry RC, Collins DL, et al. Unbiased average age-appropriate atlases for pediatric studies. Neuroimage. 2011;54(1):313–27.

134. Fonov VS, Evans AC, McKinstry RC, Almli CR, Collins DL. Unbiased nonlinear average age-appropriate brain templates from birth to adulthood. NeuroImage. 2009;47:S102.

135. Salami A, Adolfsson R, Andersson M, Blennow K, Lundquist A, Adolfsson AN, et al. Association of APOE varepsilon4 and Plasma p-tau181 with Preclinical Alzheimer’s Disease and Longitudinal Change in Hippocampus Function. J Alzheimers Dis. 2022;85(3):1309–20.

136. Janiesch C, Zschech P, Heinrich K. Machine learning and deep learning. Electronic Markets. 2021;31(3):685–95.

137. He K, Zhang X, Ren S, Sun J. Deep Residual Learning for Image Recognition. arXiv: 160806993. 2015.

138. Nair V, Hinton GE. Rectified linear units improve restricted boltzmann machines. Proceedings of the 27th International Conference on International Conference on Machine Learning; Haifa, Israel: Omnipress; 2010. p. 807–14.

139. Developers T. TensorFlow. v2.5.0 edition. https://www.tensorflow.org/ (accessed 01 June 2022).

140. Chollet F. Keras. 2.5.0 edition. https://keras.io (accessed 01 June 2022).

141. Sutskever I, Martens J, Dahl G, Hinton G. On the importance of initialization and momentum in deep learning. In: Sanjoy D, David M, editors. Proceedings of the 30th International Conference on Machine Learning; Proceedings of Machine Learning Research: PMLR; 2013. p. 1139–47.

142. Srivastava N, Hinton GE, Krizhevsky A, Sutskever I, Salakhutdinov R. Dropout: a simple way to prevent neural networks from overfitting. Journal of Machine Learning Research. 2014;15:1929–58.

143. Selvaraju RR, Cogswell M, Das A, Vedantam R, Parikh D, Batra D. Grad-CAM: Visual Explanations from Deep Networks via Gradient-Based Localization. International Journal of Computer Vision. 2020;128(2):336–59.

144. Greve DN, Salat DH, Bowen SL, Izquierdo-Garcia D, Schultz AP, Catana C, et al. Different partial volume correction methods lead to different conclusions: An (18)F-FDG-PET study of aging. Neuroimage. 2016;132:334–43.

145. Lammertsma AA, Hume SP. Simplified reference tissue model for PET receptor studies. Neuroimage. 1996;4(3 Pt 1):153–8.

146. Baron RM, Kenny DA. The moderator-mediator variable distinction in social psychological research: conceptual, strategic, and statistical considerations. J Pers Soc Psychol. 1986;51(6):1173–82.

147. Preacher KJ, Hayes AF. SPSS and SAS procedures for estimating indirect effects in simple mediation models. Behav Res Methods Instrum Comput. 2004;36(4):717–31.

148. Poldrack RA, Huckins G, Varoquaux G. Establishment of Best Practices for Evidence for Prediction: A Review. JAMA Psychiatry. 2020;77(5):534–40.

