## Supplementary materials for "Brain-Cognitive Gaps in relation to Dopamine and Health-related Factors: Insights from AI-Driven Functional Connectome Predictions"

Esmaeili et al.

The supplementary file includes Figures S1—S4 and Tables S1 & S2

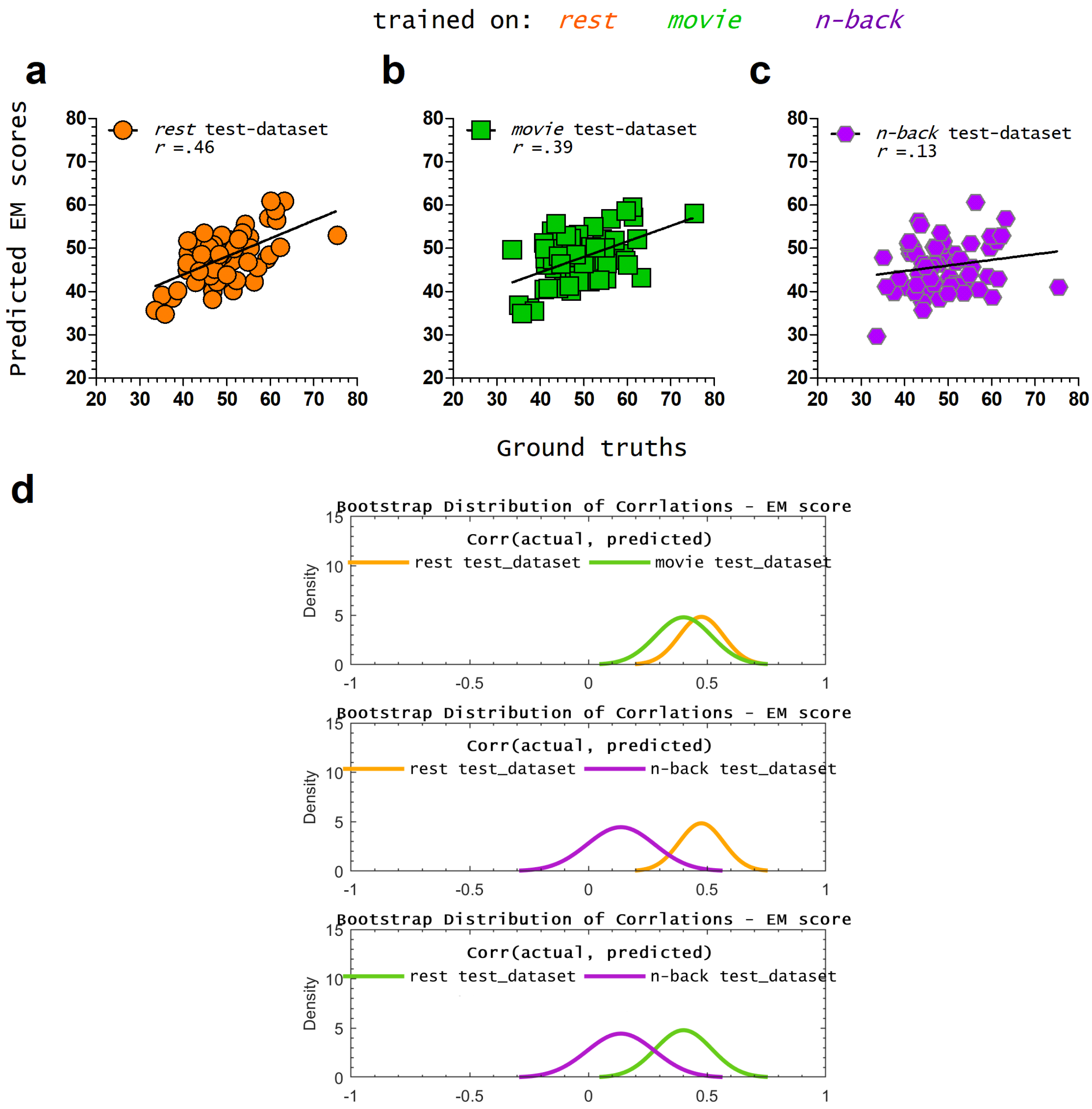


**Figure S1.** Model trained on functional connectivity maps acquired at rest predicts (**a**) episodic memory (EM) of the test dataset. The models trained on movie-watching (**b**) but not n-back (**c**) datasets predicted EM scores of the test dataset significantly. **Table S1** summarizes the *p* values and correlation power for each model. Test datasets were obtained from the same cohort for rest, movie-watching, and n-back. Bootstrap distributions of correlations between predicted and actual EM scores indicated no significant difference in the predictive power of EM between models trained on resting-state and movie-watching datasets (**d**). Additionally, the bootstrap distribution revealed that models trained on resting-state and movie-watching data yielded higher correlations than those trained on n-back data (**d**).

***Table S1.*** *Correlation results for episodic memory (EM) score predictions – DyNAMiC dataset with Schaefer300 parcellation.*

| **Model trained on** | rest | | | movie | | | n-back | | |
|---|---|---|---|---|---|---|---|---|---|
| **Model tested on** | rest | movie | n-back | rest | movie | n-back | rest | movie | n-back |
| Correlation ($r$) | **0.46** | **0.38** | **0.36** | **0.34** | **0.39** | 0.09 | 0.06 | 0.15 | 0.14 |
| Correlation ($r^2$) | **0.09** | **0.05** | **0.04** | **0.03** | **0.06** | -0.05 | -0.07 | -0.01 | -0.02 |
| Significance | **0.0002** | **0.004** | **0.005** | **.007** | **0.002** | 0.47 | 0.67 | 0.25 | 0.28 |
| MSE | 84.18 | 112.30 | 76.82 | 129.66 | 68.23 | 138.91 | 144.22 | 95.83 | 121.45 |
| MAE | 3.66 | 4.47 | 3.37 | 5.11 | 3.02 | 5.28 | 5.40 | 3.94 | 4.96 |
| COBRA | | | | | | | | | |
| Correlation | **N.A.** | | | | | | | | |
| Significance | **N.A.** | | | | | | | | |

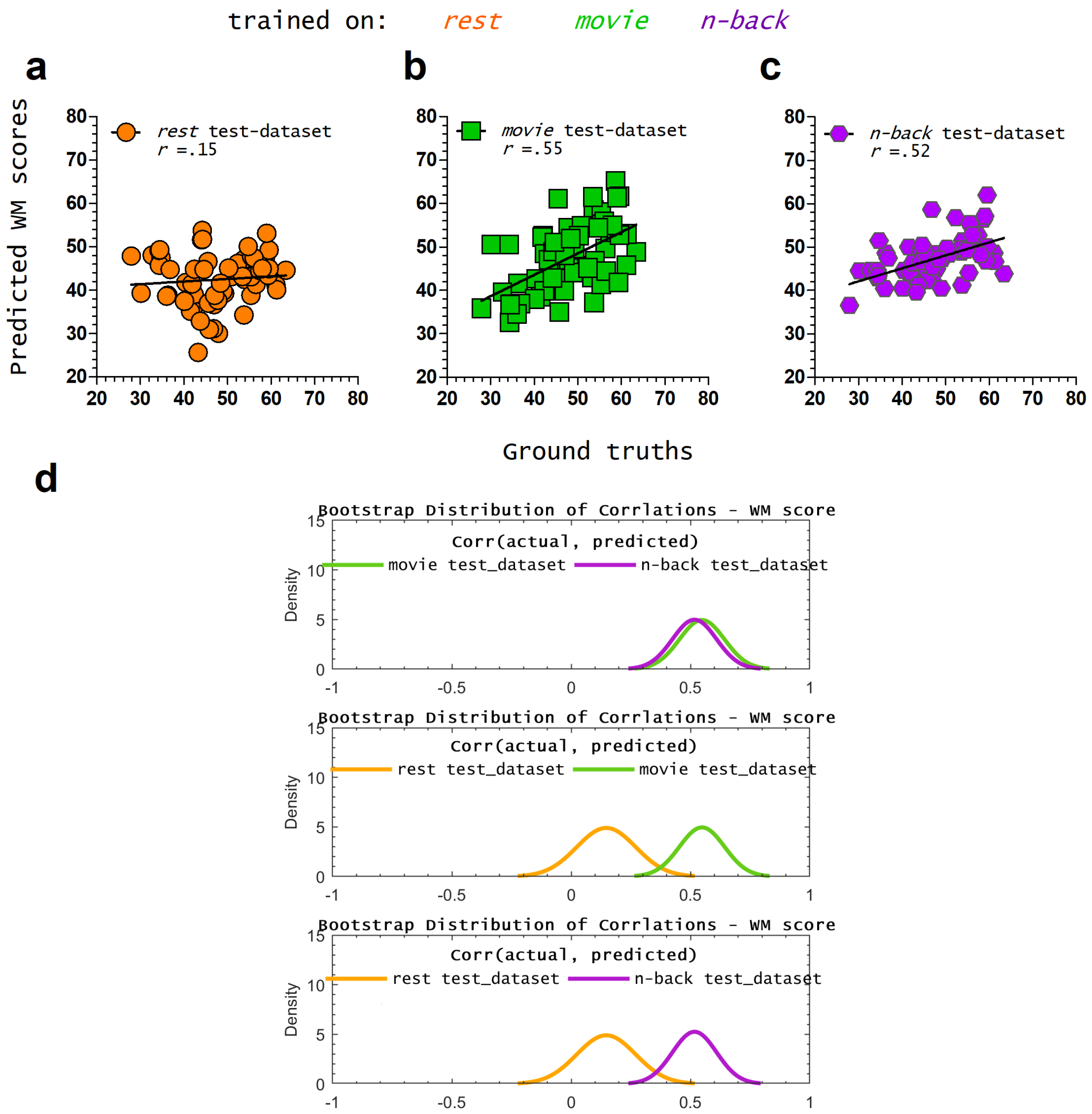


**Figure S2.** Model trained on functional connectivity maps acquired at rest did not significantly predict (**a**) the working memory (WM) of the test dataset. The corresponding model best performed in WM scores predictions while trained on the movie-watching data set (**b**). The model trained on the n-back dataset (**c**) was the second-best, predicting WM scores. **Table S2** summarizes the *p* values and correlation power for each model. Test datasets for resting-state, movie-watching, and n-back tasks were derived from the same cohort. Bootstrap distributions of correlations between predicted and actual WM scores showed no significant difference in predictive power between models trained on movie-watching and n-back data (**d**). Furthermore, the bootstrap distribution revealed that models trained on movie-watching and n-back data exhibited higher correlations than those trained on the resting state dataset (**d**).

***Table S2.*** *Correlation results for working memory (WM) score predictions – DyNAMiC dataset with Schaefer300 parcellation.*

| **Model trained on** | rest | | | movie | | | n-back | | |
|---|---|---|---|---|---|---|---|---|---|
| **Model tested on** | rest | movie | n-back | rest | movie | n-back | rest | movie | n-back |
| Correlation (*r*) | 0.16 | 0.12 | 0.10 | **0.33** | **0.55** | **0.43** | **0.38** | **0.46** | **0.52** |
| Correlation ($r^2$) | -0.01 | -0.03 | -0.04 | **0.03** | **0.16** | **0.09** | **0.05** | **0.10** | **0.14** |
| Significance | 0.27 | 0.36 | 0.43 | **0.01** | **<.0001** | **0.006** | **0.003** | **0.0002** | **<.0001** |
| MSE | 90.90 | 92.76 | 94.61 | 87.33 | 76.51 | 82.97 | 86.45 | 81.04 | 77.43 |
| MAE | 3.22 | 3.77 | 3.89 | 3.74 | 3.12 | 3.62 | 3.79 | 3.88 | 3.29 |
| COBRA | | | | | | | | | |
| Correlation | | | | | | **N.A.** | | | |
| Significance | | | | | | **N.A.** | | | |

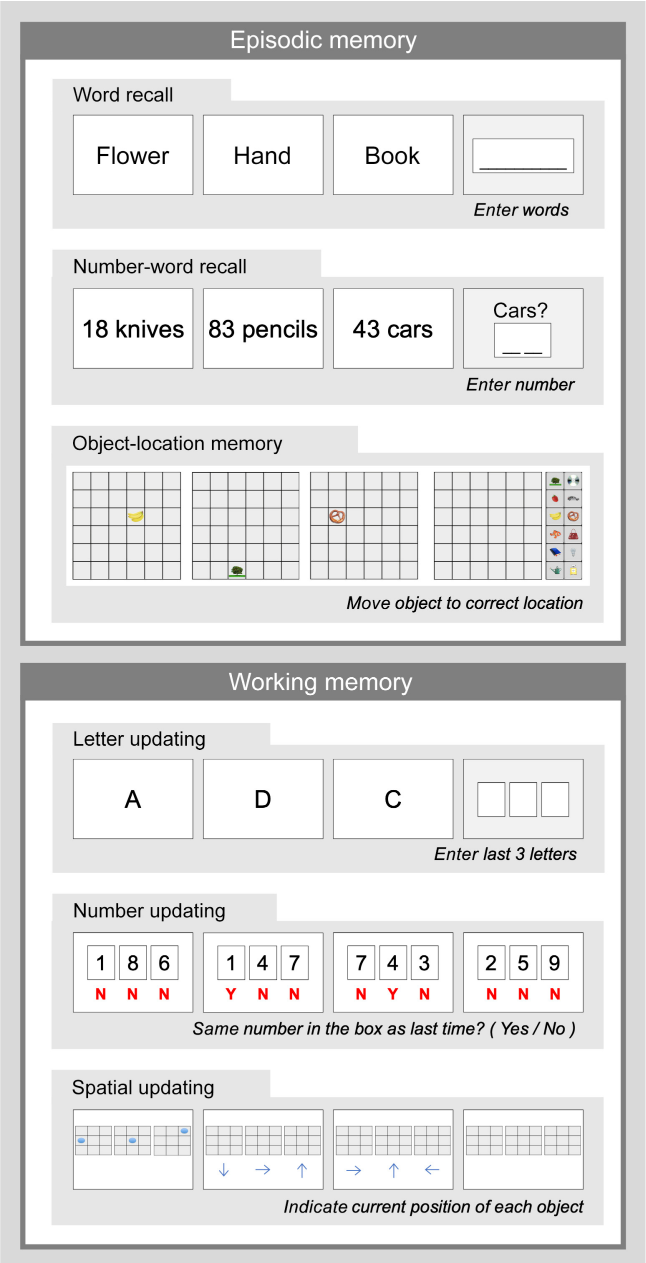


**Figure S3.** Overview of the main cognitive tests included in DyNAMiC. Illustration is adopted from Nordin et al. (1), an open-access article distributed under the terms of the Creative Commons CC BY license.

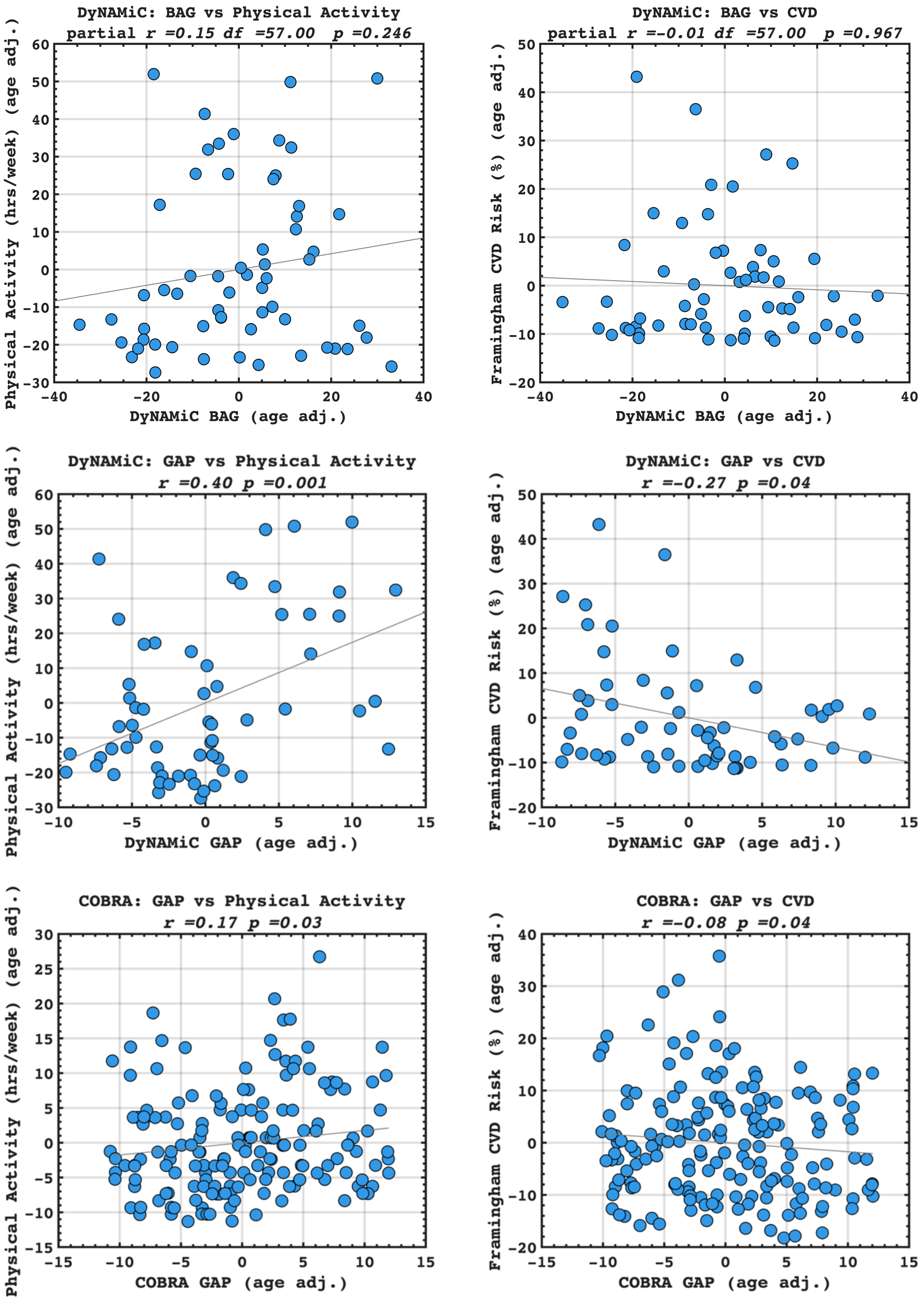


**Figure S4**. Partial correlation analysis of brain age gap (BAG) vs. physical activity and cardiovascular disease (CVD) risk (top row). Partial correlation analysis between brain-cognition gap (BCG, here referred to as GAP) vs. physical activity and CVD in DyNAMiC (middle row) and COBRA (last row) datasets.

### References

1. Nordin K, Gorbach T, Pedersen R, Panes Lundmark V, Johansson J, Andersson M, et al. DyNAMiC: A prospective longitudinal study of dopamine and brain connectomes: A new window into cognitive aging. J Neurosci Res. 2022;100(6):1296–320.
